# Cholinergic disruption of state-dependent retrosplenial layer 1 activity causes temporal associative memory deficit under stress

**DOI:** 10.1101/2025.05.27.656297

**Authors:** Asami Tanimura, Hande Login, Solbjørg Østergaard Feld-Jakobsen, Wen-Hsien Hou, Hilligje Gunnink, Chihiro Nakamoto-Griffiths, Tobias Overmark, Jelena Radulovic, Naoki Yamawaki

## Abstract

Memory functions rely on discrete patterns of neuronal activity emerging from individual neurons, their interconnected circuits, and their alignment with global brain states. We show that such patterns are generated by layer 1 inhibitory neurons in the ventral retrosplenial cortex (vRSPL1), whose activity correlated with immobility and specifically contributed to the formation of temporal associative memories. This state-dependent activity was subject to stress-induced cholinergic modulation through muscarinic 1 receptor (M1R) signaling, leading to selective impairments in temporal but not contextual associative memory. Slice studies showed that these effects were likely due to M1R-mediated inhibition of vRSPL1 neurons, transiently disrupting their local connectivity as well as responsiveness to afferent input. Together, we demonstrate a mechanism by which vRSPL1 activity aligned to immobility coordinate the formation of temporal associations. Through its sensitivity to stress-related cholinergic modulation, this mechanism presents vulnerability to traumatic amnesia, especially for temporal details of episodic memories.

## INTRODUCTION

Brain states are understood as recurring, globally distributed patterns of neuronal activity that emerge during cognitive processes and influence future behavior^1^. While functionally discrete states rely on the coordinated activity of specific brain regions, the state-dependent activity of neurons within the same region is often heterogeneous due to differences in connectivity and sensitivity to neuromodulatory tone^1^. This tone, shaped by cholinergic, aminergic, and peptidergic innervation, controls state transitions that enable rapid cognitive and behavioral adaptation to changes in internal and external environments.

Studies in rodents demonstrate that brain states can be inferred from spontaneous behavior, particularly locomotion and immobility^2^. These behaviors reflect brain states engaged in externally- or internally-directed cognition and support fundamental functions such as exploration and navigation (locomotion) or rest and sleep (immobility). Importantly, the brain states that emerge during these behaviors are linked to distinct activity patterns required for the formation and retention of episodic memories, including theta oscillations (locomotion), sharp-wave ripples (rest and sleep), and slow-wave oscillations (sleep)^3–5^.

Several lines of evidence suggest that the retrosplenial cortex (RSP) is a key locus representing memory-linked activity patterns that emerge during locomotion and immobility. During locomotion, RSP neurons conjunctively encode allocentric and egocentric information relevant for spatial navigation and learning^6–8^. During immobility, the RSP drives resting-state connectivity of the default mode network in both humans and rodents^9–11^. In humans, this activity pattern is implicated in internally-directed cognition, including memory retrieval, planning, reflection, and mind-wandering^12,13^. Yet, how state-dependent RSP functions are performed at the level of defined cell types remains unknown.

Given that the activity of individual neurons in the RSP can dynamically shift their affiliation between local and distal cortical networks depending on the animal’s locomotor state^14^, it raises the possibility that single neurons may be flexibly recruited across different states.

Alternatively, specific cell types might exhibit a preferential affiliation with particular behavioral or brain states. To address this, we investigated the activity and functions of layer 1 inhibitory neurons in the ventral retrosplenial cortex (vRSPL1), as well as their relationship with cholinergic activity associated with brain state regulation during the trace fear conditioning (TFC) paradigm. Furthermore, we explored how acute stress – a putative disruptor of neuromodulatory tone and brain state^15^ – affects these activities and functions.

Our findings reveal that vRSPL1 neurons are selectively active during immobility and play a specific role in the formation of temporal associative memories. This immobility-aligned activity is inversely correlated to the activity of cholinergic neurons in the diagonal band nucleus, which innervate vRSPL1. Moreover, the memory function of these neurons was disrupted by stress acting on their muscarinic 1 (M1) receptor, which exerted an inhibitory effect on vRSPL1 neurons, as demonstrated *ex vivo*. These results suggest that the activity of vRSPL1 neurons, aligned with immobility-linked brain states, bidirectionally regulates the formation of temporal associative memory in a manner critically dependent on neuromodulatory tone.

## RESULTS

### vRSPL1 population activity is correlated with immobility and memory processes

vRSP integrates inputs from the dorsal hippocampus and anterior thalamus, which are implicated in spatial navigation and episodic memory in layer 1^16^. Based on this, we hypothesized that neurons in this layer play a key role in mediating state-dependent functions associated with distinct behaviors. To test this, we monitored the activity of the vRSPL1 neuronal population during a TFC paradigm, which allowed us to assess vRSPL1 activity in relation to both immobility (e.g., freezing) and locomotion prior to conditioning. The main neuronal constituents of vRSPL1 were GABAergic neurons, with approximately 80% expressing the *Ndnf* gene, similar to other cortical regions^17^ (**Fig. S1**). We therefore expressed GCaMP8s in this population and recorded their activity using fiber photometry (**Fig. 1A–D**).

**Figure 1.**
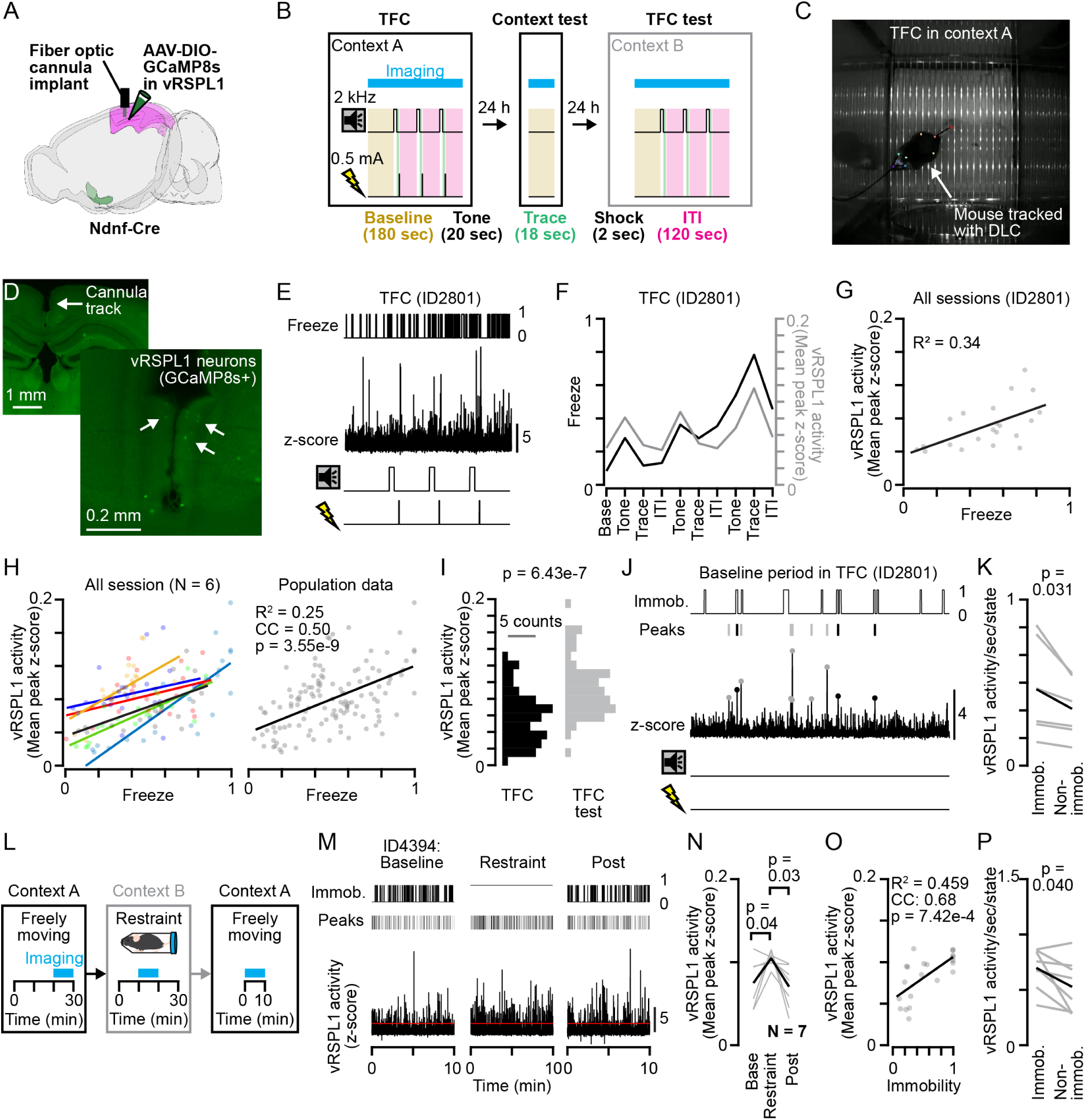
vRSPL1 population activity is correlated with the immobility and memory process. **(A)** Schematic of the surgery performed. **(B)** Schematic of the trace fear conditioning (TFC) paradigm. Timing of tone and shock presentations is represented as TTL pulses. Distinct periods of the TFC protocol (baseline, trace, and intertrial interval (ITI)) are indicated by different colors. A blue horizontal bar indicates the fiber photometry recording period. **(C)** A video frame of a mouse undergoing TFC. vRSPL1 activity is simultaneously recorded with fiber photometry. Body points tracked by DeepLabCut (DLC) are also indicated. **(D)** An example brain section image showing the track of the optic cannula, prepared after the end of the experiment. GCaMP8s-labeled neurons in vRSPL1 are also indicated with arrows. **(E)** Example traces acquired from one mouse (ID2801) during TFC. From top to bottom: freezing behavior, vRSPL1 activity, tone delivery, and shock delivery. **(F)** Freezing (black) and vRSPL1 activity (gray) across different TFC phases for mouse ID2801. **(G)** Relationship between freezing and vRSPL1 activity across TFC, context test, and TFC test in mouse ID2801. **(H)** Left: The same plot as in G but include data from other mice (N = 6, indicated with different color). Right: The same plot as the left, but regression is plotted using a population data. CC: Correlation coefficient. **(I)** Distribution of vRSPL1 activity during the TFC session (black) and the TFC test (gray) (TFC: 0.064 ± 0.030 vs TFC test: 0.098 ± 0.030; p = 6.43e-7, paired t-test, N = 6). **(J)** Example traces from the baseline period (180 sec) of (E). Black points denote calcium peaks aligned with immobility period; gray lines denote peaks not aligned with immobility. **(K)** Fraction of vRSPL1 calcium transients occurring during immobility versus non-immobility periods, normalized by the fraction of time spent in each state. Event rate during immobility: 0.461 ± 0.297 events/sec; during non-immobility: 0.342 ± 0.184 events/sec (p = 0.031, signed-rank test, N = 6). **(L)** Schematic of the protocol used to assess the effect of acute restraint stress. A blue horizontal bar indicates the fiber photometry recording period. **(M)** Example traces from one mouse (ID4394) during the restraint protocol shown in (L). From top to bottom: freezing behavior, detected vRSPL1 calcium peaks, and overall vRSPL1 activity. **(N)** vRSPL1 activity levels before, during, and after restraint. Mean peak z-score in baseline vs. restraint period: 0.075 ± 0.028 vs. 0.104 ± 0.009 (p = 0.04, paired t-test). Restraint vs. post-restraint period: 0.104 ± 0.009 vs. 0.071 ± 0.026 (p = 0.03, paired t-test). **(O)** Relationship between immobility and vRSPL1 activity across the restraint protocol, represented using a population data (N = 7 mice). **(P)** Fraction of vRSPL1 calcium transients occurring during immobility versus non-immobility periods during the baseline period in experiment (L), normalized by the fraction of time spent in each state. Event rate during immobility: 0.697 ± 0.178 events/sec; during non-immobility: 0.527 ± 0.240 events/sec (p = 0.04, paired t-test, N = 7).

During the TFC (we used ‘TFC’ to describe the conditioning phase of the TFC paradigm; **Fig. 1B**), vRSPL1 activity gradually increased with repeated presentations of the tone–trace– shock sequence, closely paralleling the rise in freezing behavior (**Fig. 1E,F, Methods**). When we plotted population activity against freezing across different phases of TFC and memory tests, we observed a strong correlation between the two measures (**Fig. 1G,H**). A direct comparison of vRSPL1 activity between the TFC and TFC test revealed higher activity during the TFC test (**Fig. 1I**).

Next, we examined vRSPL1 activity during the baseline period of the TFC. This analysis revealed that vRSPL1 activity occurred more frequently during the period of immobility than locomotion (**Fig. 1J,K**).

Given the preferential engagement of the vRSPL1 population during freezing or immobility, we asked whether this response generalizes to externally imposed immobility. To test this, we measured vRSPL1 activity before, during, and after physical restraint, which triggers stress responses^18^ (**Fig. 1L**). vRSPL1 activity increased during restraint and returned to baseline level after restraint (**Fig.1M,N**). Plotting vRSPL1 activity against the immobility for each phase from all animals revealed a positive correlation (**Fig. 1O**). Consistent with our earlier findings, vRSPL1 activity during the baseline period in this experiment also occurred more frequently during spontaneous immobility (**Fig. 1P**).

We then asked whether vRSPL1 activity precedes or follows the onset of the tone, footshock, or immobility by aligning their activity to each event onset. This analysis revealed that vRSPL1 activity did not follow the tone (or precisely, it did not respond to a neutral tone) but increased following the onset of either footshock or immobility (**Fig. S2A–F, M,N**). The vRSPL1 activity is decoupled to immobility during the footshock, when their activity coincided with short movement burst (**Fig. S2C,D**)

Together, these data suggest that vRSPL1 activity is primarily aligned to a brain state linked to immobility and cognitive processes related to associative learning, associative memories, and stress.

### DBN^Chat^ population activity is anti-correlated with freezing and immobility

We next investigated what controls the shift of vRSPL1 activity during the TFC and memory tests. Given that such state transitions are often regulated by neuromodulators^1,2,19^, we examined the potential contribution of cholinergic neurons in the diagonal band nucleus (DBN^Chat^), which preferentially projected to layer 1 of the vRSP regardless of its subregions (**Fig. S3**). To assess their role, we expressed GCaMP8s in DBN^Chat^ neurons and monitored their population activity during the TFC and memory tests (**Fig. 2A,B, Methods**). In contrast to the vRSPL1 population, DBN^Chat^ activity was anti-correlated with freezing behavior (**Fig. 2C–E**). A direct comparison of DBN^Chat^ population activity between the TFC and TFC test revealed higher activity during the TFC (**Fig. 2F**).

**Figure 2.**
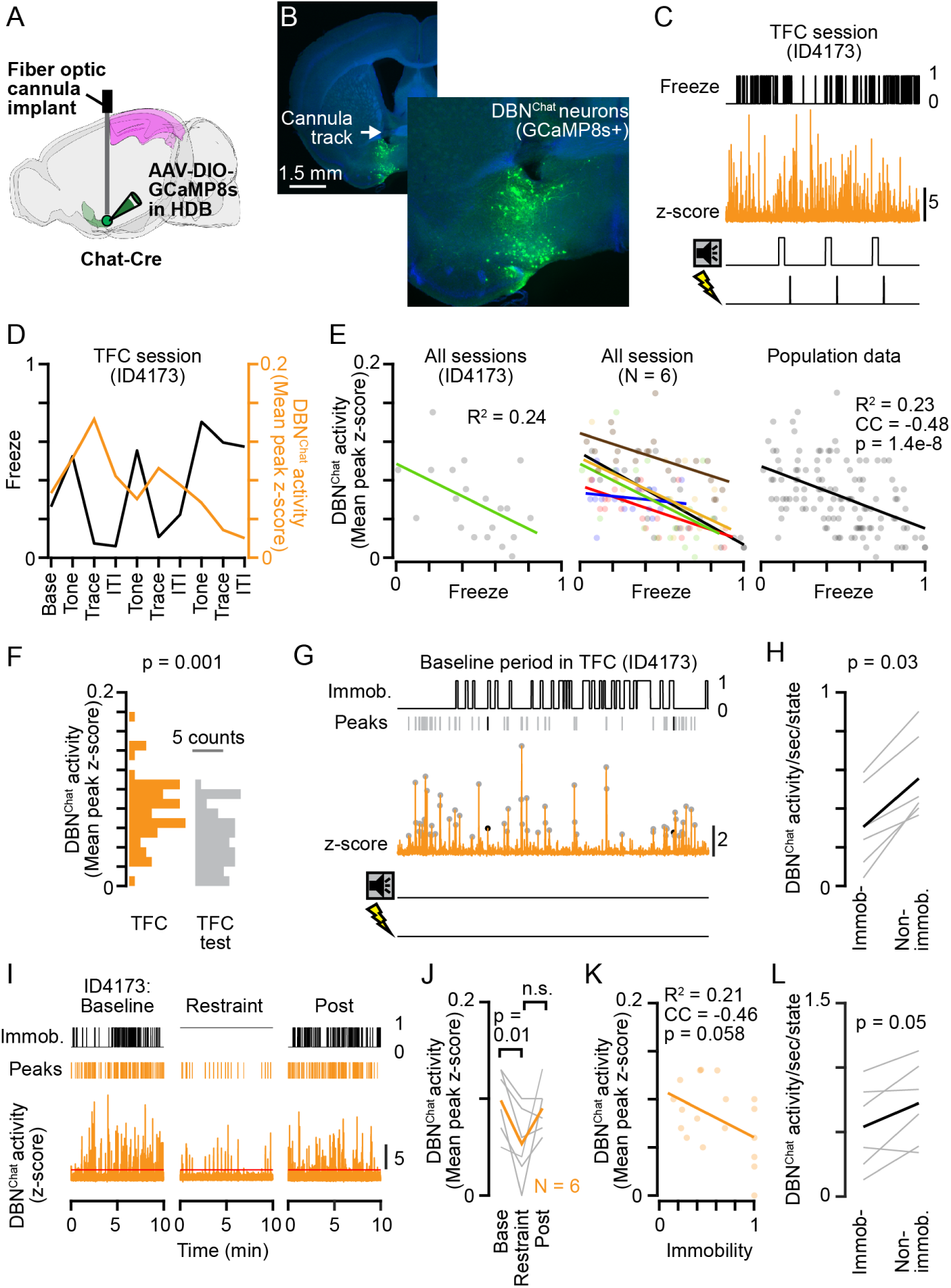
DBN^Chat^ population activity is anti-correlated with freezing and immobility. **(A)** Schematic of the surgery performed. **(B)** An example brain section image showing the track of the optic cannula (arrow) and GCaMP8s-labeled neurons in DBN. **(C)** Example traces acquired from one mouse (ID4173) during TFC. From top to bottom: freezing behavior, DBN^Chat^ activity, tone delivery, and shock delivery. **(D)** Freezing (black) and DBN^Chat^ activity (orange) across different TFC phases for mouse ID4173. **(E)** Relationship between freezing and DBN^Chat^ activity across the TFC, context test, and TFC test. Left: Data from a single mouse (ID4173). Middle: Data from additional mice (N = 6), each shown in a different color. Right: Linear fit using a population data from all mice. CC: Correlation coefficient. **(F)** Distribution of DBN^Chat^ activity during the TFC session (yellow) and the TFC test (gray) (TFC: 0.075 ± 0.034 vs TFC test: 0.055 ± 0.030; p = 6.43e-7, paired t-test, N = 6). **(G)** Example traces from the baseline period (180 sec) of (E). Black points denote calcium peaks aligned with immobility period; gray lines denote peaks not aligned with immobility. **(H)** Fraction of DBN^Chat^ calcium transients occurring during immobility versus non-immobility periods, normalized by the fraction of time spent in each state. Event rate during immobility: 0.305 ± 0.218 events/sec; during non-immobility: 0.556 ± 0.224 events/sec (p = 0.031, signed-rank test, N = 6). **(I)** Example traces from one mouse (ID4173) during the restraint protocol. From top to bottom: freezing behavior, detected DBN^Chat^ calcium peaks, and overall DBN^Chat^ activity. **(J)** DBN^Chat^ activity levels before, during, and after restraint. Mean peak z-score in baseline vs. restraint period: 0.098 ± 0.034 vs. 0.053 ± 0.035 (p = 0.01, paired t-test). Restraint vs. post-restraint period: 0.053 ± 0.035 vs. 0.089 ± 0.025 (p = 0.12, paired t-test). **(K)** Relationship between immobility and DBN^Chat^ activity across the restraint protocol, represented using population data. **(L)** The same as (H), but from data obtained in restraint experiment. Event rate during immobility: 0.541 ± 0.336 events/sec; during non-immobility: 0.725 ± 0.322 events/sec (p = 0.05, signed-rank test).

To determine whether DBN^Chat^ activity was specifically anti-correlated with freezing or more generally with an immobility, we analyzed their activity during the baseline period of the TFC. We found that DBN^Chat^ activity occurred more frequently during a period of non-immobility (**Fig. 2G,H**).

We further examine this relationship under externally imposed immobility, a manipulation previously linked to elevate cholinergic tone^20,21^. Consistent with our findings made in the TFC experiment, DBN^Chat^ activity was reduced during restraint compared to baseline (**Fig. 2I,J**). No difference was observed between activity levels during the restraint and post-restraint periods in this experiment, likely due to increased immobility in some animals following restraint (**Fig. 2J,K**). Plotting DBN^Chat^ activity against immobility confirmed an anti-correlated relationship, and DBN^Chat^ activity at baseline occurred more frequently during a period of non-immobility in this experiment (**Fig. 2K,L**). Moreover, the alignment of DBN^Chat^ activity to immobility onset revealed a decrease in their activity following the transition to immobility (**S2G,H,O,P**), mirroring the vRSPL1 activity changes but in the opposite direction.

These findings indicate that the activity of DBN^Chat^ neurons, which project to vRSPL1, is primarily aligned with a brain state associated with locomotion—a state opposite to the one with which vRSPL1 neurons are aligned.

### Stress or inhibition of vRSPL1 activity impairs temporal associative memory formation

We next assessed how memory processes and activity of the vRSPL1 population are affected by stress by applying restraint stress prior to TFC (**Fig. 3A,B**). This manipulation did not alter baseline activity or freezing behavior during the TFC on average, but it reduced freezing during both context and TFC tests (**Fig. 3C-E**). When vRSPL1 activity was plotted against freezing behavior, a positive correlation remained (**Fig. 3F-H**). Moreover, vRSPL1 activity remained higher during the TFC test (**Fig. 3I**). These findings suggest that vRSPL1 population activity continues to align with freezing and memory processes after stress.

**Figure 3.**
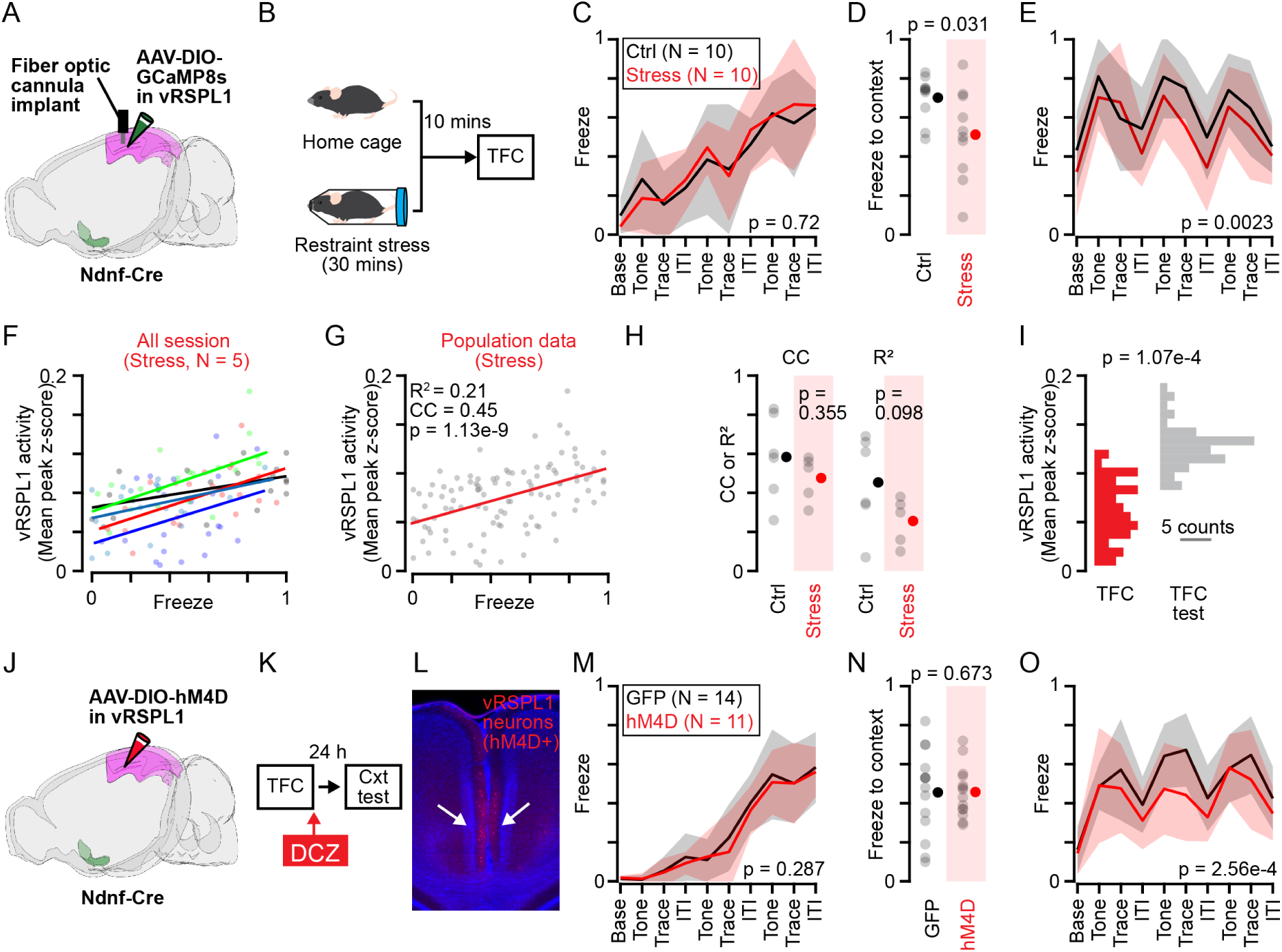
Stress or inhibition of vRSPL1 activity impairs temporal associative memory formation. **(A)** Schematic of the surgery performed. **(B)** Schematic of the experiment performed, highlighting the timing of restraint stress relative to TFC paradigms. **(C)** Freezing during TFC for control mice (black) and stressed mice (red). Two-way ANOVA: main effect of phase, F_(9,180)_ = 19.65, p = 1.02e-22; effect of group, F_(1,180)_ = 0.13, p = 0.722; interaction between phase and group, F_(9,130)_ = 0.43, p = 0.916. **(D)** Freezing during context test for control and stress mice (p = 0.031, rank-sum test). **(E)** Freezing during TFC test for control and stress mice. Two-way ANOVA: main effect of phase, F_(9,180)_ = 9.08, p = 2.80e-11; effect of group, F_(1,180)_ = 9.57, p = 0.0023; interaction between phase and group, F_(9,130)_ = 0.59, p = 0.797. **(F)** Relationship between freezing and vRSPL1 activity across TFC, context test, and TFC test in stressed mice (N = 5, different mice indicated with different color). **(G)** The same as (F), but regression is plotted using a population data. **(H)** Correlation coefficient (CC) and R² obtained from control and stressed mice (CC control: 0.58 ± 0.22 vs. CC stress: 0.48 ± 0.12, p = 0.355, unpaired t-test; R² control: 0.45 ± 0.24 vs. R² stress: 0.23 ± 0.10, p = 0.098, unpaired t-test). **(I)** Distribution of vRSPL1 activity during the TFC session (red) and the TFC test (gray) in stressed mice (TFC: 0.062 ± 0.033 vs TFC test: 0.090 ± 0.027; p = 1.07e-4, signed-rank test). **(J)** Schematic of the surgery performed. **(K)** Schematic of the experiment performed, highlighting the timing of DCZ injection relative to TFC paradigms. **(L)** Epifluorescence image showing vRSPL1 neurons expressing hM4D (indicated with arrows). **(M)** Freezing during TFC for control mice (black) and hM4D mice (red). Two-way ANOVA: main effect of phase, F_(9,130)_ = 64.67, p = 4.73e-58; effect of group, F_(1,130)_ = 1.14, p = 0.287; interaction between phase and group, F_(9,130)_ = 0.24, p = 0.988. **(N)** Freezing during context test for control and hM4D mice (p = 0.673, unpaired t-test). **(O)** Freezing during TFC test for control and hM4D mice. Two-way ANOVA: main effect of phase, F_(9,130)_ = 13.45, p = 2.89e-17; effect of group, F_(1,130)_ = 13.80, p = 2.56e-4; interaction between phase and group, F_(9,130)_ = 0.99, p = 0.447.

The freezing deficit observed during the context and TFC tests but not during the TFC suggests that stress disrupts the brain state necessary for memory formation during the post-TFC period. Given the high correlation of vRSPL1 activity with memory-triggered freezing behavior (**Fig. 1**), we hypothesized that stress disrupted memory-related vRSPL1 activity during post-TFC period and caused memory impairment. To test this, we selectively inhibited vRSPL1 neurons after TFC using hM4D(Gi) in non-stressed mice, activated via intraperitoneal administration of the selective agonist deschloroclozapine (DCZ) (**Fig. 3J–L**). This manipulation reduced freezing during the TFC test without affecting freezing in the context test (**Fig. 3M-O**). DCZ administration in mice lacking hM4D(Gi) expression had no effect on baseline movement or freezing behavior (**Fig. S4**).

Together, these findings demonstrate that restraint stress prior to TFC impairs memory by disrupting post-learning brain states, a period during which vRSPL1 activity is essential specifically for temporal associative memory formation.

### Selective knockdown of muscarinic 1 (M1) receptors in vRSPL1 neurons rescues the stress-induced temporal associative memory deficit

The anti-correlated relationship between DBN^Chat^ and vRSPL1 activity in relation to freezing behavior, along with the impact of vRSPL1 activity inhibition on the TFC test, suggests that cholinergic inhibition of vRSPL1 neurons could be a mechanism contributing to stress-induced memory deficits. Muscarinic receptors, in particular, have been implicated in brain state transitions associated with movement and cholinergic tone^22,23^. We therefore investigated whether muscarinic receptors expressed in vRSPL1 neurons mediate the effects of stress on memory.

To explore this, we first examined the expression of muscarinic receptor subtypes in vRSPL1 neurons. Transcriptomic data from the Allen Brain Atlas revealed that cortical layer 1 neurons, primarily from primary visual area and identified by the gene markers *Ndnf*, *α7nAChR*, and *VIP*, express both muscarinic M1 and M3 receptors (**Fig. 4A**). Our RNAscope analysis confirmed the expression of M1 receptors (M1R) in vRSPL1 neurons, but showed minimal to no expression of M3 receptors (**Fig. 4B,C**).

**Figure 4.**
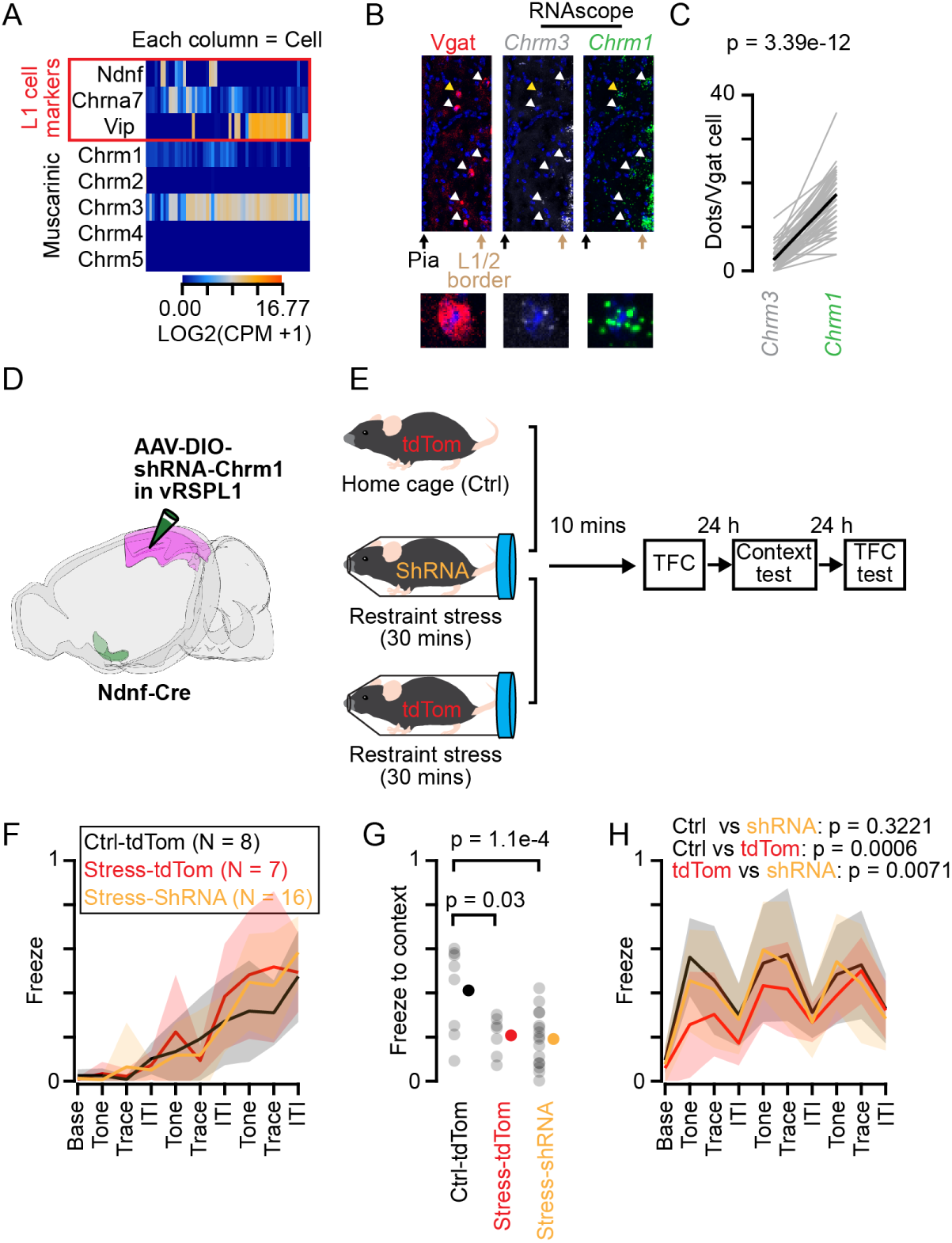
Selective knockdown of muscarinic 1 (M1) receptors in vRSPL1 neurons rescues the stress-induced temporal associative memory deficit. **(A)** Heatmap showing mRNA expression levels of muscarinic receptor subtypes (*Chrm1*– *Chrm5*) in layer 1 cortical neurons derived from Allen Brain Atlas scRNA-seq database. **(B)** Confocal image of vRSPL1 showing *Vgat^+^* somata (red) co-labeled with RNAScope probes for *Chrm1* (green) and *Chrm3* (gray). **(C)** Transcript abundance quantified by counting puncta per cell using a z-projected image (*Chrm3*: 2.0 ± 2.1 puncta per cell. *Chrm1*: 13.9 ± 5.7 puncta per cell. p = 4.13e-07 (signed-rank test). n = 65 cells, N = 3 mice). **(D)** Schematic of the surgery performed. **(E)** Schematic of the experiment performed. **(F)** Comparison of freezing between non-stressed mice with tdTomato-expression (black), stressed mice with tdTomato expression (red) and stressed mice with shRNA-expression (yellow) during TFC. Two-way ANOVA. For control vs stress-tdTom mice: main effect of phase, F_(9,130)_ = 16.95, p = 3.14e-18, effect of group, F_(1,130)_ = 2.86, p = 0.0932; interaction, F_(9,130)_ = 1.15, p = 0.3358. For control vs stress-shRNA mice: main effect of phase, F_(9,220)_ = 28.39, p = 2.26e-32, effect of group, F_(1,220)_ = 1.24, p = 0.267; interaction, F_(9,220)_ = 1.22, p = 0.283. For stress-tdTom vs stress-shRNA mice: main effect of phase, F_(9,210)_ = 29.55, p = 7.14e-33, effect of group, F_(1,210)_ = 0.92, p = 0.340; interaction, F_(9,210)_ = 0.76, p = 0.658. **(G)** Freezing during context test between three groups (For control vs stress-tdTom mice, p = 0.03 (unpaired t-test). For control vs stress-shRNA mice, p = 1.1e-4 (unpaired t-test). For stress-tdTom vs stress-shRNA mice, p = 0.680 (unpaired t-test)). **(H)** Freezing during TFC test between three groups. Two-way ANOVA. For control vs stress-tdTom mice: main effect of phase, F_(9,130)_ = 8.11, p = 1.72e-9, effect of group, F_(1,130)_ = 12.24, p = 6.0e-4; interaction, F_(9,130)_ = 0.850, p = 0.571. For control vs stress-shRNA mice: effect of phase, F_(9,220)_ = 11.48, p = 1.01e-14, effect of group, F_(1,220)_ = 0.98, p = 0.322; interaction, F_(9,220)_ = 0.34, p = 0.959. For stress-tdTom vs stress-shRNA mice: effects of phase, F_(9,210)_ = 9.6, p = 3.10e-12; effect of group, F_(1,210)_ = 7.38, p = 0.0071, interaction, F_(9,210)_ = 1.0, p = 0.444.

To test if stress-induced memory deficit is caused by M1R-dependent modulation of vRSPL1 activity, we selectively knocked down the receptor by targeting *Chrm1* mRNA using a Cre-dependent shRNA virus (**Fig. 4D, Methods**). Control experiments confirmed that this manipulation reduced *Chrm1* mRNA expression by approximately 80% in vRSPL1 neurons, without significantly affecting *Chrna7* mRNA levels (**Fig. S5**). We then assigned mice to three experimental groups: (1) a control group injected with a Cre-dependent tdTomato virus in vRSPL1 with no exposure to stress, (2) a stress group injected with the same control virus and subjected to restraint stress, and (3) a stress group injected with the *Chrm1* shRNA virus and subjected to restraint stress (**Fig. 4E**). All mice were subsequently subjected to TFC, followed by context test and TFC test.

Relative to the control group, both stressed tdTomato- and shRNA-injected mice exhibited intact acquisition but reduced freezing during the context test, consistent with our earlier findings (**Fig. 4F,G**). However, a divergence emerged during the TFC test: while the stressed tdTomato group continued to show reduced freezing compared to controls, the stressed shRNA group exhibited freezing behavior comparable to that of the non-stressed control group (**Fig. 4H**).

These results suggest that stress impairs temporal associative memory by promoting cholinergic signaling, which disrupts vRSPL1 activity through an M1R-dependent mechanism. Combined with findings from the hM4D(Gi)-mediated inhibition experiment, these data further support a specific role for vRSPL1 neurons in regulating temporal, but not contextual, associative memory formation.

### Cholinergic signaling reduces the excitability of vRSPL1 neurons and reduces its responsiveness to afferent input

To investigate how cholinergic signaling from DBN^Chat^ neurons modulates vRSPL1 activity, and thereby contributes to its state dependency, we performed whole-cell recordings from vRSPL1 neurons *ex vivo* and examined the effects of the cholinergic receptor agonist carbachol (CCh, 1 µM) on their passive and active membrane properties (**Fig. 5A**). On average, CCh reduced the input resistance of vRSPL1 neurons, while having minimal effect on resting membrane potential (**Fig. S6A,B**). Moreover, CCh reduced the neurons’ responsiveness to injected currents, producing a rightward shift in the input-output (f-I) curve (**Fig. 5B,C; Fig. S6A-D**).

**Figure 5.**
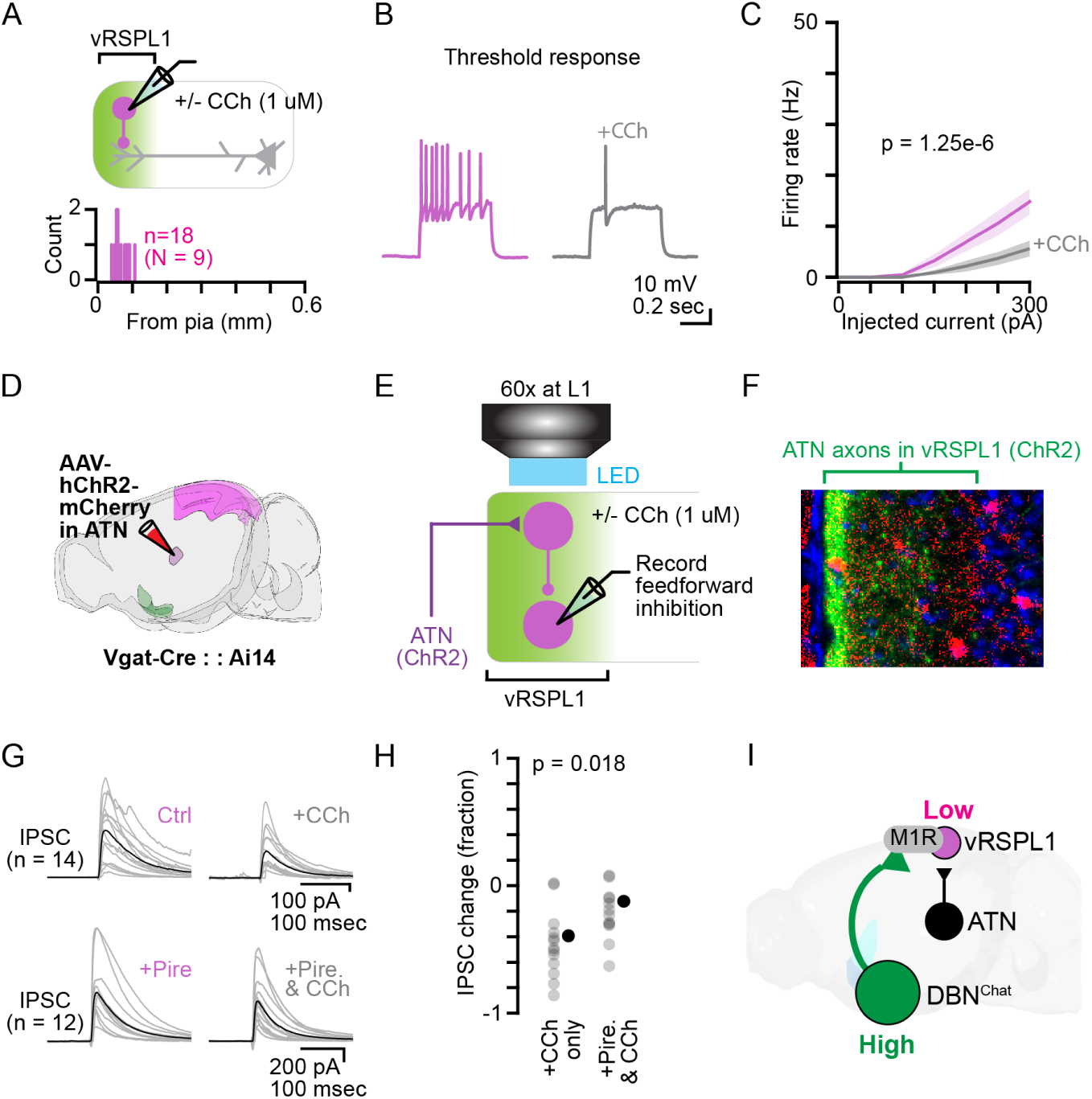
Cholinergic signaling reduces the excitability of vRSPL1 neurons and reduces its responsiveness to afferent input. **(A)** Top: Schematic of ex vivo whole-cell recordings from vRSPL1 neurons in acute brain slices. Bottom: Histogram showing the distribution of cortical depths of recorded neurons. **(B)** Representative spike traces from a vRSPL1 neuron in response to threshold current injection before (magenta) and after (gray) bath application of carbachol (CCh). **(C)** Input-output (f-I) curve of vRSPL1 neurons before (magenta) and after (gray) CCh application. Two-way ANOVA: main effect of current, F_(6,238)_ = 20.12, p = 5.29e-19; effect of CCh, F_(1,238)_ = 24.75, p = 1.25e-6; interaction between current and CCh, F_(6,238)_ = 4.19, p = 5.01e-4; n = 18 cells from 9 mice. **(D)** Schematic of the surgical procedure used to express ChR2 in thalamic axons and label GABAergic neurons in vRSPL1. **(E)** Schematic of the circuit being tested and ex vivo whole-cell recordings performed to isolate feedforward inhibition. **(F)** Top: Epifluorescence image showing vRSPL1 with ChR2-expressing ATN axons (green), tdTomato-labeled *Vgat^+^*neurons (red), and DAPI (blue). Bottom: Histogram displaying the distribution of cortical depths of recorded vRSPL1 neurons. **(G)** Top: Photo-evoked IPSCs recorded in vRSPL1 neurons in response to ATN terminal stimulation, before and after CCh application. Each gray trace represents an individual neuron; black trace indicates the average response across neurons (control: 470.8 ± 305.0 pA vs CCh: 232.9 ± 194.2 pA, p = 0.001 (signed-rank test); n = 13 cells from 8 mice). Bottom: Same experiment performed in the presence of the M1R antagonist pirenzepine (control: 607.6 ± 415.5 pA vs CCh: 459.3 ± 315.6 pA, p = 0.002 (signed-rank test); n = 12 cells from 6 mice). **(H)** Fraction of change in photo-evoked IPSC by CCh with or without pirenzepine (CCh only: -0.46 ± 0.26 vs Pire and CCh: -0.22 ± 0.20 pA, p = 0.018 (unpaired t-test)). **(I)** A model of the circuit interaction between vRSPL1 and DBN^Chat^ neurons based on our study.

For comparison, we examined the effects of CCh on pyramidal neurons in layers 5A and 5B. CCh had no effect on layer 5A neurons (**Fig. S6E-H**) but depolarized the resting membrane potential and enhanced excitability in layer 5B neurons (**Fig. S6I-L**).

To determine if these effects are mediated by M1R, we performed f-I curve analyses under various pharmacological conditions (**Fig. S7**). The CCh-induced shift in the curve was blocked by co-application of scopolamine (1 µM) and mecamylamine (10 µM), a generic muscarinic and nicotinic receptor antagonist, or by scopolamine alone, suggesting muscarinic receptor involvement. Mecamylamine alone did not prevent the effect, further implicating muscarinic rather than nicotinic receptors. The application of the M1R antagonist pirenzepine (1 µM) partially prevented the shift, indicating that M1R mediates the cholinergic suppression of vRSPL1 excitability, but not all.

The reduced excitability of vRSPL1 neurons by CCh suggests that these neurons become less responsive to afferent inputs under this condition. To identify the afferent sources of input to vRSPL1 neurons, we first performed monosynaptic rabies tracing, which revealed multiple presynaptic regions, including the DBN, anterior thalamic nuclei (ATN), and dorsal hippocampus (including neurons in both the subiculum and stratum radiatum–stratum lacunosum-moleculare border of CA1) (**Fig. S8A–D)**. To assess how cholinergic signaling modulates afferent recruitment of vRSPL1 neurons, we focused on an excitatory pathway from ATN that projects predominantly to upper layer 1 but forms uniform connections with all vRSPL1 neurons, as determined in our connectivity study (**Fig. S8E–I**).

We expressed ChR2 in ATN axons and performed whole-cell recordings from vRSPL1 neurons to measure feedforward inhibition evoked by optogenetic stimulation of ATN axons before and after CCh application (**Fig. 5D-H**). We found photo-evoked feedforward inhibition was reduced by ∼40% after CCh application (**Fig. 5G,H**). Similar effects were observed when feedforward inhibition was measured from layer 5B neurons (**Fig. S9A-D**).

To further complement these findings, we assessed the effect of CCh on inhibitory input from vRSPL1 neurons to local pyramidal neurons by expressing soma-targeted ChR2 (ChroME^24^) in vRSPL1 neurons, chosen to facilitate somatic action potential-dependent GABA release, rather than release induced by direct excitation of axon terminals. Recordings from layer 5B neurons showed that CCh application reduced the inhibitory input to local pyramidal neurons (**Fig. S9E–H**).

To determine whether these effects were mediated by M1R, we measured feedforward inhibition onto vRSPL1 neurons in the presence of the pirenzepine. Pirenzepine attenuated the CCh-induced reduction in feedforward inhibition approximately by half (**Fig. 5G,H**), indicating the major involvement of M1R in this attenuation.

Together, these findings suggest that cholinergic signaling decreases the excitability of vRSPL1 neurons via M1R-dependent mechanism, thereby by making them less responsive to L1-targeting afferent inputs (**Fig. 5I**).

## DISCUSSION

By combining genetic, molecular, and circuit-level tools with behavioral paradigms, we identified cell-type-specific, state-dependent, and stress-sensitive mechanisms underlying temporal associative memory formation. These mechanisms are mediated by vRSPL1 neurons, which exhibit elevated activity during immobility and are highly sensitive to stress-induced cholinergic modulation through M1R (**Fig. 6**).

**Figure 6.**
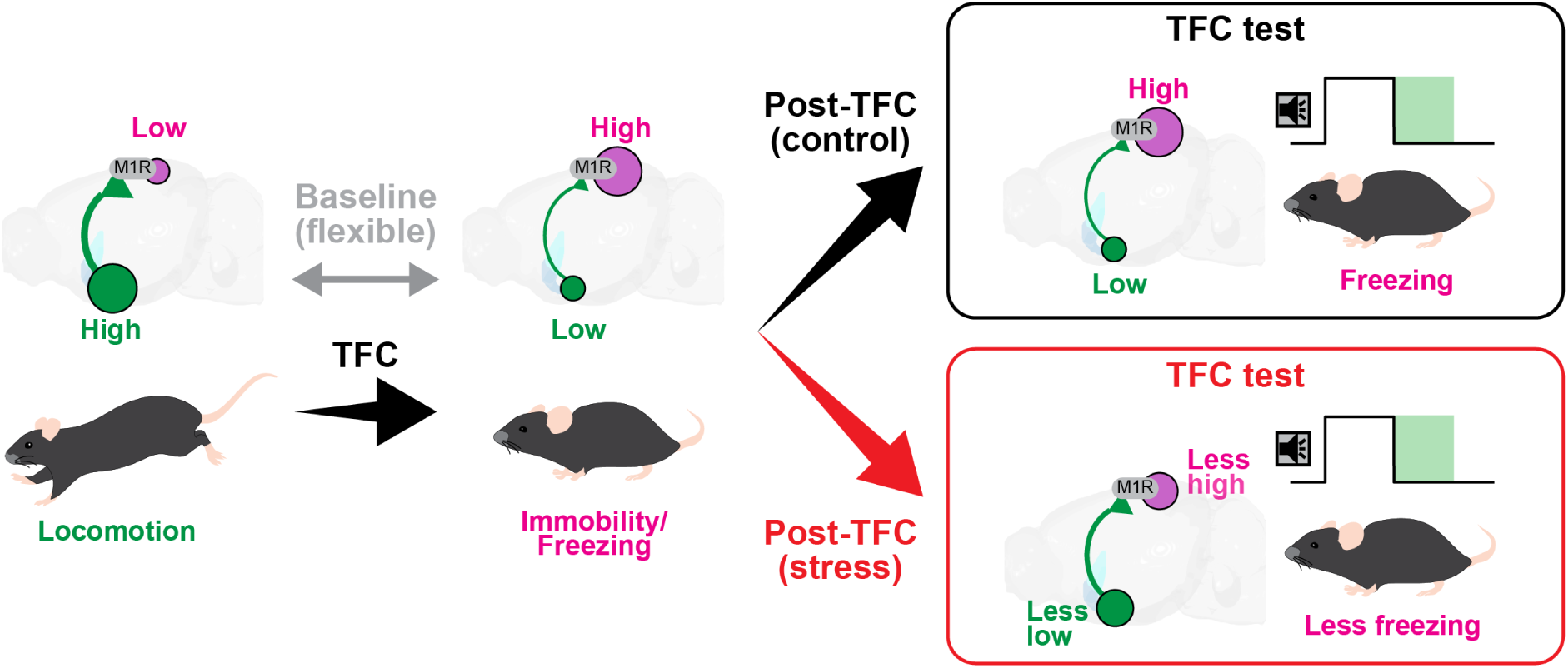
A model of stress effect on brain states aligned with vRSPL1 activity and temporal associative memory. There are two opposing brain states: one dominated by DBNChat activity (green), linked to locomotion and externally directed cognition; and another dominated by vRSPL1 activity (magenta), linked to immobility and internally directed cognition. These states flexibly alternate at baseline (gray arrow), but TFC biases the system toward the vRSPL1-dominant state (black arrow). Stress does not affect this state shift during TFC, but it alters vRSPL1 activity during the post-TFC period via dysregulated activation of M1R (red arrow). This induces a specific deficit in temporal associative memory formation without affecting contextual associative memories.

This distinguishes vRSPL1 neurons from layer 1 neurons identified by *Ndnf* in primary sensory cortices, which exhibit increased activity during locomotion^25^, or encode memory by increasing responses to conditioned stimuli^26,27^. While some differences may reflect methodological variation (e.g., single-cell versus population imaging), they more likely arise from the unique connectivity of vRSPL1 with the episodic memory network. Specifically, vRSPL1 is one of the few cortical regions that receive direct, monosynaptic input from both the dorsal hippocampus and ATN^16,28^, which encode episodic and spatial memory. Our rabies tracing and slice electrophysiology experiments confirmed these monosynaptic connections, supporting the idea that, rather than responding to external sensory input, vRSPL1 neurons predominantly respond to internal information during immobility. An exception to the internally driven nature of vRSPL1 activity occurred during footshock, which evoked a brief burst of movement. Footshocks recruit multiple neural circuits beyond primary sensory pathways^29^ and has been shown to activate layer 1 neurons via cholinergic input acting on nicotinic receptors^27^. Thus, a similar mechanism may underlie the footshock-evoked responses observed in vRSPL1 neurons.

In contrast to vRSPL1 neurons, DBN^Chat^ neurons, which also project to vRSPL1, exhibited an opposite activity pattern during freezing and immobility. This aligns with prior studies showing that basal forebrain cholinergic neurons, including those in the DBN, encode movement and locomotion^22,30–34^. Because the correlation to locomotion occurs at the level of somata, axons, and acetylcholine release^35^, the somatic activity we observed in DBN^Chat^ neurons likely reflects acetylcholine dynamics at their projection sites, including vRSPL1. A likely consequence of sustained cholinergic input during locomotion is the desynchronization of network activity^36^, which is predicted to favor the processing of external sensory inputs by the network^37^, contrasting with the predominant processing of internal information by vRSPL1 neurons in absence of locomotion.

Although vRSPL1 activity was elevated during various forms of immobility, its emergence during the acquisition and later retention of temporal associative memory was particularly notable as it was tightly linked to the immobility induced by tone-shock pairings. This type of immobility (freezing) differs from spontaneous immobility in that it co-occurs with other emerging brain states, such as attention and arousal, governed by neuromodulatory circuits.

The generation of such complex brain states may synchronize distinct neuronal populations aligned to those states for the rapid formation and long-term retention of selected memories^2^, such as memories of negative experiences.

Importantly, while vRSPL1 activity was necessary for temporal associative memory and its modulation by stress, it had no effect on contextual associative memory under any testing condition. This distinction occurred despite the conditions being matched for negative value and brain states. Given that the formation and persistence of contextual associative memory in the vRSP primarily depends on L2/3 neurons receiving excitatory input from the subiculum^38,39^, these findings highlight a functional segregation within vRSP microcircuits for different forms of episodic memory.

One potential circuit supporting the distinct function of vRSPL1 neurons is their connection to long-range GABAergic projections from CA1 neurons at the stratum radiatum–stratum lacunosum-moleculare border, as shown in our data ^16,40,41^. These CA1 neurons are likely downstream of entorhinal “island cells,” which are critical for temporal associative memory^42,43^. Thus, vRSPL1 may serve as a specialized cortical hub for consolidating temporal information processed by the island cell → CA1 pathway.

Knockdown of M1Rs in vRSPL1 neurons selectively rescued the stress-induced deficit in temporal associative memory, further supporting our findings indicating their specialized role in this process. This also implicates cholinergic signaling as a mediator of stress-induced disruption in vRSPL1 activity. Notably, M1R knockdown had no effect on acquisition, suggesting that M1Rs in vRSPL1 are not involved in acquisition per se, but primarily target post-learning memory formation.

Our *ex vivo* experiments demonstrate that prolonged cholinergic signaling inhibits vRSPL1 neurons by reducing input resistance via M1Rs, providing a mechanistic basis for the state-dependent activity of these neurons observed in vivo. However, our data also indicated a contribution from other cholinergic receptors, perhaps, M3, to the effect, as pirenzepine was unable to fully block the CCh effects. Muscarinic receptors desensitize far more slowly (∼20 minutes^44,45^) than nicotinic receptors (∼100 ms^46,47^), and cholinergic axons exhibit sustained activity during locomotion^22,30^, suggesting that muscarinic signaling likely predominates in this behavioral state. The most plausible explanation for the M1R effect is the activation of calcium-dependent SK channels via the Gq–IP3 signaling cascade^48,49^. This M1R-dependent reduction in excitability renders vRSPL1 neurons less responsive to afferent input, as demonstrated in our study. While this effect may facilitate local network desynchronization during locomotion (e.g., via disinhibition), it could also contribute to associative memory deficits if dysregulated—potentially explaining the stress effect observed in our study. These results also highlight that M1Rs expressed on vRSPL1 neurons as a promising therapeutic target for disorders involving disrupted temporal processing, such as stress-induced temporal amnesia observed in post-traumatic stress disorder.

What remains unresolved is what this emergent activity represents and what drives it. While it clearly reflects cognitive processes within the TFC paradigm, its function during spontaneous immobility or restraint stress remains unclear. One potential clue lies in its resemblance to patterns observed in the default-mode network and sharp-wave ripples (SWRs), both of which emerge during immobility and are associated with internally-directed cognitive functions – such as planning, memory retrieval, and memory consolidation – in rodents^3,5,50,51^, as well as reflection, imagination, and mind-wandering in humans^12,13^. Thus, vRSPL1 activity during distinct immobile states may reflect engagement in one of these internal processes. Indeed, its involvement in memory consolidation and retrieval is strongly suggested by our data, and elucidating the dynamics of vRSPL1 activity during the post-TFC period remains a key goal for future studies. Some of the vRSPL1 activity we observed may be driven by SWRs propagated from dorsal CA1, which is linked to memory consolidation^5^. This is consistent with previous findings showing that the RSP is a main cortical region activated shortly after SWR peaks in dorsal CA1^39,52^. Notably, this activation precedes or coincides with suppressed subcortical activity, including in the thalamus^53,54^, supporting the idea that dorsal CA1 is a key driver of vRSPL1 activity. However, thalamocortical inputs may also contribute to vRSPL1 dynamics, particularly in support of working memory and sensory integration during learning or memory acquisition^55,56^. Future experiments will be needed to disentangle the relative contributions of different afferent inputs to vRSPL1 activity across distinct behavioral states.

Taken together, we propose that the vRSPL1 neuronal population serves as a state-dependent, stress-sensitive cortical coordinator of hippocampal–neocortical temporal memory representation, organizing temporal associations within local retrosplenial networks through its extensive inhibitory connectivity.

## METHODS

All experiments were conducted in accordance with standard ethical guidelines and were approved by the Danish National Animal Experiment Committee (License number: 2021-15-0201-00801). Unless otherwise specified, all animals used were transgenic mice with a C57BL/6J background. Mice were 2–4 months old at the time of experiment. We did not select for a specific sex and used both male and female mice as they became available in our colony. Animals were housed under a 12-hour light/dark cycle (lights on at 6:00 a.m.).

The transgenic mouse lines used in this study included:

- B6.Cg-Ndnf^tm1.1(folA/cre)Hze/J^ (*Ndnf*-Cre, JAX#028536)
- B6J.129S6(FVB)-Slc32a1^tm2(cre)Lowl^/MwarJ (*Vgat*-Cre, JAX#028862)
- B6;129S6-ChAT^tm2(cre)Lowl^/J (*Chat*-Cre, JAX#006410)

In some experiments, these lines were crossed with B6.Cg-Gt(ROSA)26Sor^tm^^14(CAG-tdTomato)Hze^/J (Ai14, JAX#007914) mice, and offspring from these crosses were used for analysis.

### Viruses

The information on viruses used in our experiments is the following.

- AAV5-hEF1a-dlox-EGFP(rev)-dlox (v217-5, VVF)
- AAV1-hSyn1-dlox-jGCaMP8s(rev) (v627-1, VVF)
- AAV8-hSyn-DIO-hM4Di (Gi) – mCherry (44362-AAV8, Addgene)
- AAV5-CAG-GFP (37525-AAV5, Addgene)
- AAV5-CAG.hChR2(H134R)-mCherry (10054-AAV5, Addgene)
- AAV5-EF1a-DIO-hChR2(E123T/T159C)-EYFP (35509-AAV5, Addgene)
- AAV1-hSyn1-dlox-TVA_2A_mCherry_2A_oG(rev)_dlox (v306-1, VVF)
- Rabies-GFP (NTNU viral core facility)
- AAV9-CAG-DIO-ChroME-ST-P2A-H2B-mRuby3 (108912-AAV9, Addgene)
- AAV9-FLEX-Syn1-mCherry-shRNA-*Chrm1* (custom ordered , VectorBuilder)

### Stereotaxic injection

Mice were anesthetized with isoflurane (3 % for induction, 1.5–2 % for maintenance) and placed in a stereotaxic frame (Model 940, Kopf Instruments) equipped with a feedback-controlled heating pad (ThermoStar Homeothermic System, RWD) set to maintain body temperature at 38 °C. To prevent corneal drying, ophthalmic gel was applied to the eyes. Preemptive analgesia was administered subcutaneously using buprenorphine (0.1 mg/kg) and meloxicam (1.5 mg/kg).

The scalp was sterilized using isopropyl alcohol, and a midline incision was made to expose the skull. Bregma and lambda were identified and aligned to ensure proper skull leveling.

Craniotomies were performed using 0.5 mm drill bits (Fine Science Tools). Stereotaxic coordinates for target brain regions were as follows (in mm):

- vRSPL1: AP: −1.90 and −2.10; ML: 0.10 (0.00 for cannula); DV: 0.75 and 0.90
- DBN: AP: +1.00; ML: 1.00; DV: 5.20 and 5.50
- ATN: AP: −0.85; ML: 1.10; DV: 3.25

For virus delivery, beveled glass pipettes (Wiretrol II, Drummond Scientific) front-filled with viral solution were lowered to the target coordinates. Injections of 50–100 nL were performed using a custom-built displacement-based injector (based on the MO-10, Narishige). After injection, the pipette was left in place for 3 minutes before slow retraction to minimize backflow.

Postoperative care included daily subcutaneous injections of meloxicam for 3 days. Mice were allowed to recover and were used for experiments 3–5 weeks post-injection.

### Optic cannula implantation

For fiber photometry recordings, fiber optic cannula (1.25 mm ceramic ferrule, 400 µm core diameter, 0.39 NA, 1.0 or 5.5 mm fiber length) were implanted targeting either the vRSPL1 or horizontal DBN. Cannula implantation was performed either on the same day as viral injection or approximately two weeks later, using a custom cannula holder for precise placement.

To prepare the skull, overlying connective tissue was removed using 0.3% hydrogen peroxide, and the skull surface was gently scored with a scalpel blade to improve adhesion. Once the cannula was positioned at the stereotaxically defined coordinates, any exposed brain surface around the fiber was sealed using Kwik-Cast (World Precision Instruments). The cannula was then secured to the skull with dental adhesive cement (Super-Bond, SUN MEDICAL). After curing, the entire skull surface was coated with carboxylate cement (Durelon, 3M ESPE) for added stability and protection.

### Fiber photometry

The fiber photometry system (all components from Doric Lenses) consisted of a console, LED driver, and a cube integrated with U.V. and blue LEDs (iiFMC5). The cube was connected to the mice’s optical cannula via a rotary joint (FRJ_1x1 FC-FC) and two patch cords (core 400 um, 0.37NA, low auto-fluorescent). GCaMP8s was excited with blue LED (470 nm), and the isosbestic point was excited with U.V. LED (415 nm). The power for each wavelength was ∼35 µW and ∼15 µW, respectively, measured at the tip and calibrated before each experiment on the day of recording. Hardware control and data acquisition were managed by Doric Neuroscience Studio software (DNS, V6). The behavior recording and fiber photometry were synchronized by triggering DNS and Ethovision with a common TTL pulse delivered via an Arduino running a custom-written program. The photometry signals were acquired in lock-in mode, and post-processed using the python and MATLAB functions (see below).

### Acute restraint stress combined with trace fear conditioning

Three to five days before the behavioral tests, mice were single-housed and kept in the Scantainer located in Room B, adjacent to the behavior rooms (Rooms A and C). Two to three days before the experiment, mice were habituated to handling and fiber photometry recording by attaching the patch cord for 5 minutes daily.

On the day of the experiment, mice were randomly assigned to either the control or stress group. The control group underwent trace fear conditioning (TFC) in Room C directly from their home cage. In contrast, the stress group was exposed to 30 minutes of restraint stress in Room A using a custom-fabricated restraint tube (modified from a 50-mL Falcon tube), followed by a 10-minute recovery period in their home cage in Room B, before undergoing TFC in Room C.

In the TFC paradigm, mice were placed in Context A, located inside a soundproof chamber equipped with a speaker. The context consisted of a square-shaped box with a metal grid floor and a 70% ethanol scent, introduced by cleaning the box approximately 5 minutes before the mice entered. The dimensions of the box were either 25.5 × 25.5 cm (for standard experiments) or 17 × 17 cm (for experiments involving fiber photometry). After 180 seconds baseline, a pure tone (2 kHz, 70 dB) was delivered for 20 seconds, followed by an 18-second trace period and a 2-second footshock (0.5 mA). This sequence of events (tone, trace, and shock) was repeated two more times, with an inter-trial interval (ITI) of 120 seconds. After the final footshock, mice remained in the box for an additional 120 seconds before being returned to their home cage.

Approximately 24 hours later, mice underwent a context test, which involved re-exposing them to Context A for 180 seconds. Another ∼24 hours later, mice underwent a TFC test, which involved exposing them to Context B. Context B had the same dimensions as Context A but featured a different wall and floor texture, as well as a distinct scent (30% bleach in water). In this test, mice were exposed to the same sequence of events as in the conditioning phase, except no footshock was delivered. The total trial duration for the TFC test was identical to that of the conditioning phase. Both the TFC and memory tests were performed in darkness (i.e. no visible light inside the box).

For experiments involving deschloroclozapine (DCZ^57^, Hello Bio), the solution was freshly prepared on the day of the experiment by dissolving DCZ powder in sterile saline (0.01 mg/10 mL) and administered at a dose of 0.01 mg/kg. Only mice with confirmed expression of hM4D in the vRSPL1 were included in the data analysis.

For experiments involving shRNA, we conducted two sessions: one comparing stress-tdTom and stress-shRNA, and another comparing control-tdTom and stress-shRNA. Behavioral analysis revealed no significant differences in the stress-shRNA group between the two sessions (p > 0.05). Therefore, data from the stress-shRNA group were pooled across sessions for subsequent analyses.

The hardware control, including video recording, was managed using EthoVision software (Noldus), which was triggered by a TTL pulse sent from the Arduino board. Video recording was made at 25 fps. The data shown in Fig. 1 (control) and Fig. 3 (stress) were collected as paired groups from the same experimental sessions.

### Recording neuronal activity before, during, and after acute restraint stress

In a subset of experiments, we examined the effect of acute restraint stress on population activity by recording it before, during, and after the restraint. After 2-3 days of handling and acclimatization to the patch cord, mice were placed in an open arena (20.5 × 26.5 cm) in Room A for 30 minutes, and behavior and neuronal activity was measured in its last 10 minutes. Next, in Room C, mice were placed into a custom-designed restraint tube that could accommodate the patch cord attachment (with a narrow slit at the top through which the cannula could be inserted) and left there for 30 minutes. Activity was monitored during the middle 10 minutes of the restraint period. After the restraint period, mice were released and allowed to rest in their home cage in Room B for 10 minutes before being returned to the same arena in Room A where they had been before the restraint. Activity was then measured again. Video acquisition was performed using a camera (model DMK 37AUX273, The Imaging Source) controlled by the Bonsai program at 25 fps, which was simultaneously triggered with DNS photometry signal acquisition via Arduino.

### Behavior quantification and fiber photometry data analysis

All behaviors were tracked using DeepLabCut (DLC)^58^ with a network trained on ∼1000 video frames of mice in TFC contexts A and B. The model used nine body markers: nose, both ears, head (implant), neck, both legs, tail base, and tail midpoint. The resulting time-series data for each point were processed using BehaviorDEPOT^59^, which classified freezing behavior on a frame-by-frame basis according to the following thresholds: velocity = 0.52, angle = 12°, window width = 32 frames, count threshold = 10, and minimum duration = 0.9 seconds. The key output was a binary vector indicating the freezing status in each frame, where 1 denoted freezing and 0 denoted non-freezing. This classifier and thresholds were applied consistently across all analysis. The terms ‘freeze’ and ‘non-freeze’ are used in reference to immobile behavior after conditioning, while ‘immobile’ and ‘non-immobile’ describe behavior before conditioning. A co-product of the freezing vector in BehaviorDEPOT, *movement (cm/sec)*, was used to assess baseline activity in control mice injected with either DCZ or saline (**Fig. S4**).

We did not track the mice during the restraint period in **Fig. 1L** and **Fig. 2I**, as our DLC configuration did not reliably capture behavior across all videos. Since restraint largely enforced immobility (except for occasional ‘wiggling’), we assigned a value of ‘1’ to represent behavior during this period in both experiments.

Fiber photometry signals, consisting of both calcium-dependent and calcium-independent fluorescence channels, were pre-processed prior to analysis. First, both voltage signals were low-pass filtered using a 40 Hz Butterworth filter to reduce high-frequency noise. Baseline correction was then performed using the asymmetrically reweighted penalized least squares (arPLS) algorithm. To normalize the data, z-scores were calculated for each signal. A linear regression was then performed by plotting the calcium-dependent signal against the calcium-independent signal to model baseline drift, which was subsequently subtracted from the calcium-dependent trace to isolate the activity-dependent component. To compare population activity across sessions (e.g., during TFC and memory tests) within the same animal, z-score was calculated using the global mean and standard deviation calculated from all arPLS-corrected voltage signals across sessions.

To align neural activity with behavioral data, the z-scored calcium traces were down-sampled to 25 Hz without phase shift to match the behavioral sampling rate. Peak detection was then performed on the down-sampled signal, with peaks defined as local maxima exceeding a z-score of 1.95 (corresponding to α = 0.05). For baseline event analysis, a more stringent threshold was applied, using the first trough in the peak amplitude histogram (smoothed with kernel density estimate) to isolate large events. These detected events were used to calculate mean peak amplitude and event frequency, either per second or within specific experimental phases (e.g., baseline, tone, trace, shock, and ITI). Freezing fraction was computed for each corresponding time bin using the same nesting structure.

In the plot showing freezing behavior during the TFC or TFC test, the shock period was omitted because the behavior was consistently extreme (freeze = 0), and to allow better visualization of behavioral changes during the other phases.

### *Ex vivo* electrophysiology

Mice were deeply anesthetized with isoflurane and decapitated. Coronal sections (300 µm) containing the RSP were prepared using a vibratome (VT1200S, Leica) in ice-cold choline solution, which contained (in mM): 25 NaHCO3, 1.25 NaH2PO4-H2O, 2.5 KCl, 0.5 CaCl2, 7 MgCl2, 25 D-glucose, 110 Choline chloride, 11.6 Ascorbic acid, and 3.1 C3H3NaO3. Slices were incubated in artificial cerebrospinal fluid (ACSF) containing (in mM): 125 NaCl, 25 NaHCO3, 1.25 NaH2PO4-H2O, 2.5 KCl, 11 D-glucose, 2 CaCl2, and 1 MgCl at ∼34°C for 30 minutes and then at room temperature (∼20°C) for at least 1 hour before recording.

Whole-cell recordings were performed with an upright microscope (BX51WI, Olympus) equipped with a motor-controlled stage and focus (MP285A and MPC-200, Sutter Instrument), differential interference contrast, coolLED (pE300), and a monochrome camera (Moment, Teledyne). Neurons were visualized using a 60x lens (1.00 NA, LUMPlanFL, Olympus) with Micro-Manager-2.0 gamma (NIH software). Pipettes (4-5 MΩ) were pulled from thick-walled borosilicate capillary glass (P-1000, Sutter Instrument). For voltage-clamp recordings, the internal solution contained (in mM): 135 CsMeSO_3_, 10 HEPES, 4 Mg-ATP, 0.3 Na-GTP, 8 Na_2_-Phosphocreatine, and 3.3 QX-314 (pH 7.35 adjusted with CsOH). For current-clamp recordings, CsMeSO_3,_ Na_2_-Phosphocreatine, and QX-314 was replaced with 120 KMeSO3, 8 NaCl, and 10 KCl, and pH was adjusted with KOH. Recordings and hardware control were managed with Wavesurfer (Janelia Farm). Signals were amplified and Bessel-filtered at 4 kHz with a MultiClamp 700B amplifier (Molecular Devices), then sampled at 10 kHz (connectivity) or 40 kHz (intrinsics) with a data acquisition board (USB-6343, National Instruments). Recordings with access resistance changes > 20% from baseline (∼30 MΩ) were discarded. Liquid junction potential was not corrected. All recordings were performed at ∼34°C using an inline heating system (TC-324C, Warner Instruments).

To stimulate ChR2, blue LED pulses (5 msec duration) were delivered through a minimized aperture using a 60× objective lens positioned over layer 1. In a few subset, a 4× objective (0.16 NA, UPlanSApo, Olympus) was used. The light power at the focal plane was ∼60 µW for the 60× lens and ∼20 mW for the 4× lens (corresponding to the “5%” setting on the CoolLED system). Photostimulation was repeated 3–5 times at intervals of 10 or 20 seconds to obtain averaged response traces. Mean response amplitudes were calculated within a 50 msec window following LED onset. All current-clamp recording was made in I=0 mode in Multiclamp amplifier. All pharmacological agents were purchased from Merck, TOCRIS, or HelloBio.

### Immunohistochemistry and cell type identification in vRSPL1

*Vgat*::Ai14 and *Ndnf::*Ai14 mice were transcardially perfused with ice-cold phosphate-buffered saline (PBS), followed by 4% paraformaldehyde (PFA) in PBS. Brains were dissected, post-fixed in 4% PFA at 4 °C for 24 hours, then cryoprotected sequentially in 20% and 30% sucrose in PBS at 4 °C. Tissue was embedded in Tissue-Tek O.C.T. compound (Sakura) and stored at −80 °C. Prior to sectioning, blocks were equilibrated to −20 °C, and 100-μm-thick coronal sections were collected using a cryostat (Leica CM3050 S).

Sections were incubated in blocking buffer containing 5% normal goat serum and 0.2% Triton X-100 in PBS for 30 minutes at room temperature, followed by overnight incubation at 4 °C with rabbit anti-NeuN primary antibody (1:1000; ABN78, Millipore). After PBS washes, sections were incubated for 1 hour at room temperature with Alexa Fluor 488-conjugated goat anti-rabbit secondary antibodies (1:500; A11008 or A11036, Thermo Fisher). Sections were washed again with PBS, counterstained with DAPI, and mounted using DAKO mounting medium (S3023, Agilent Technologies).

Single-plane images were acquired using a Zeiss confocal microscope equipped with a 10× objective (1024 × 1024 pixel resolution). For each brain, 2–4 images were collected across RSP sections (between AP -1.9 to -2.2 mm). Cell numbers were quantified from each image and averaged per brain. Data shown in Fig. S1 represent quantification from N = 3 brains analyzed using this method.

### Quantifying cholinergic receptor genes with RNAscope

RNAscope was used to visualize and quantify *Chrm1*, *Chrm3*, and *Chrna7* transcript expression in the vRSPL1 of *Vgat*::Ai14 mice, and to verify *Chrm1* knockdown in the same region of *Ndnf*-Cre mice injected with a Cre-dependent shRNA-*Chrm1*-mCherry viral construct.

Following fixation in 4% PFA and cryoprotection in sucrose, 20 μm-thick coronal sections containing the RSP were prepared and mounted onto Superfrost Plus slides (Fisher Scientific, 12-550-15). Slides were air-dried at −20 °C for 2 hours and stored at −80 °C.

RNAscope was performed according to the manufacturer’s protocol. To detect *Chrm1*, *Chrm3*, *Chrna7* mRNA and mCherry protein, the RNAscope™ Multiplex Fluorescent Reagent Kit v2 (ACD Biotechne, #323100) and the RNA-Protein Co-Detection Ancillary Kit (ACD Biotechne, #323180) were used. Slides were first dried at 60 °C for 30 minutes, then sequentially dehydrated in 50%, 75%, and two 100% ethanol incubations (5 minutes each), treated with hydrogen peroxide for 10 minutes at room temperature, and boiled in co-detection Target Retrieval Reagent (∼98 °C) for 5 minutes. After rinsing, sections were incubated overnight at 4 °C with anti-mCherry antibody (1:1000, Abcam ab205402) diluted in co-detection antibody diluent.

The next day, sections were post-fixed in 10% neutral buffered formalin (VWR, GEN0786-1056), incubated with Protease Plus, and rinsed with sterile water. Hybridization was performed at 40 °C for 2 hours with the following probes: *Mm-Chrm1-C1* (ACD Biotechne, 495291), *Mm-Chrm3-C2* (437701-C2), *Mm-Chrna7-C3* (465161-C3), *Mm-PPIB* (positive control; 320881), and *dabB* (negative control; 320871). Sections were stored overnight in 5× saline-sodium citrate buffer.

Signal amplification was carried out with AMP 1 (30 min), AMP 2 (30 min), and AMP 3 (15 min), all at 40 °C, with two 2-minute washes in ACD wash buffer between each step.

Fluorescent detection was performed using TSA Vivid Fluorophore Kits: Kit 650 (Biotechne, 7527) for Channel 1 (C1) and Kit 520 (Biotechne, 7523) for Channels 2 and 3 (C2, C3). After completing the ISH HRP blocking step, sections were incubated for 30 minutes at room temperature with Cy™3-conjugated donkey anti-chicken IgY secondary antibody (1:300, Jackson ImmunoResearch, 703-165-155) diluted in co-detection antibody diluent. DAPI staining was then performed, and sections were mounted using ProLong Gold Antifade Mountant (Thermo Fisher Scientific, #33342).

Imaging was conducted using a Zeiss LSM 780 confocal microscope equipped with Zen Black software. Both a 20× air objective and a 63× water-immersion objective were used (1024 × 1024 pixel resolution), with laser excitation at 405, 488, 561, and 638 nm.

For transcript quantification, images acquired with the 63× objective were analyzed using *QuPath* v0.4.3 (Bankhead et al., 2017). Regions of interest (ROIs) were manually drawn around mCherry-positive cell bodies. “Cell Detection” was applied using DAPI nuclear staining, followed by the “Subcellular Detection” tool to count individual transcript puncta per cell for each target probe. Two sections per mouse were analyzed.

### Monosynaptic rabies tracing of vRSPL1 neurons

Rabies tracing was performed as previously described^60^. Briefly, following stereotaxic injection, *Ndnf-*Cre mice were anesthetized with ketamine (120 mg/kg) and xylazine (24 mg/kg), then transcardially perfused with 4% PFA. Brains were post-fixed in 4% PFA at 4 °C, cryoprotected in 20% and then 30% sucrose in PBS, embedded in O.C.T., and frozen. Coronal brain sections (100 μm thick) were prepared using a cryostat, rinsed in PBS, counterstained with DAPI, and mounted on glass slides with coverslips using mounting medium.

Starter cells and presynaptic neuron distributions were visualized from epifluorescence images acquired on a slidescanner (Olympus VS120, 10× objective) and analyzed using AMaSiNe. For quantitative analysis, only regions containing more than 1% of the total presynaptic population were included. The fraction of presynaptic cells in each region was calculated by dividing the regional presynaptic count by the total number of labeled presynaptic neurons.

### Statistics

All data analyses were performed using MATLAB R2021b (MathWorks). Group data are presented as arithmetic means, with variability indicated by either standard deviation or individual data points, as specified. Statistical comparisons were conducted using either parametric or non-parametric tests, depending on the outcome of the normality test (Lilliefors test, **Table 1**). For parametric tests, two-tailed Student’s t-tests (paired or unpaired) were used. Otherwise, the Wilcoxon signed-rank test was applied for paired samples, and the Wilcoxon rank-sum test (Mann–Whitney U test) for unpaired samples. For comparisons involving multiple groups, two-way or N-way ANOVA was performed, followed by Scheffé’s post hoc test when significant effects were detected (**Table 2, 3 and 4**). Pearson correlation coefficients were calculated to assess the strength and direction of linear relationships between variables. Statistical significance was defined as p ≤ 0.05. The number of animals is reported as “N” and the number of neurons as “n”.

**Table 1.**
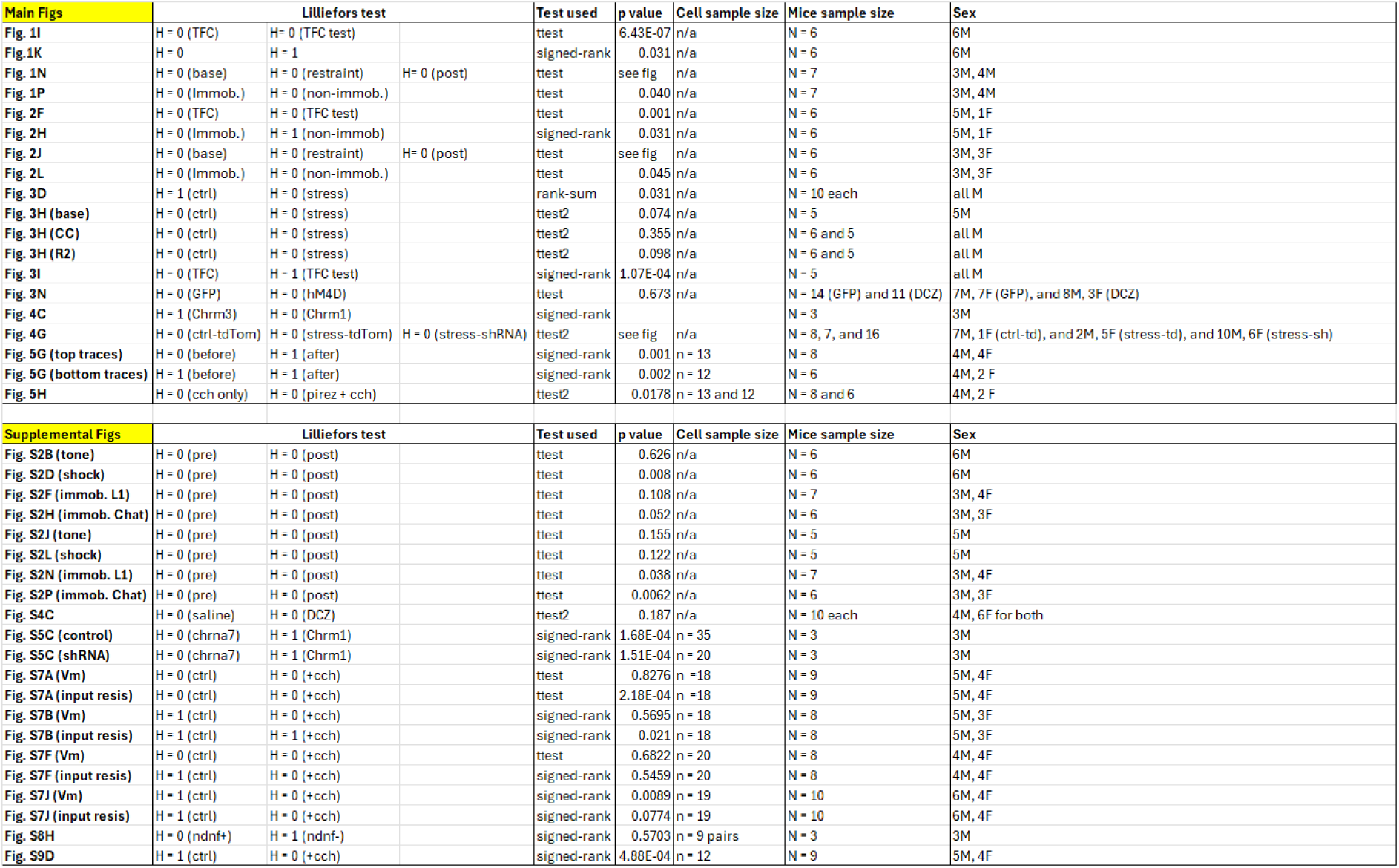
Summary of statistical tests used for paired comparisons.

**Table 2.**
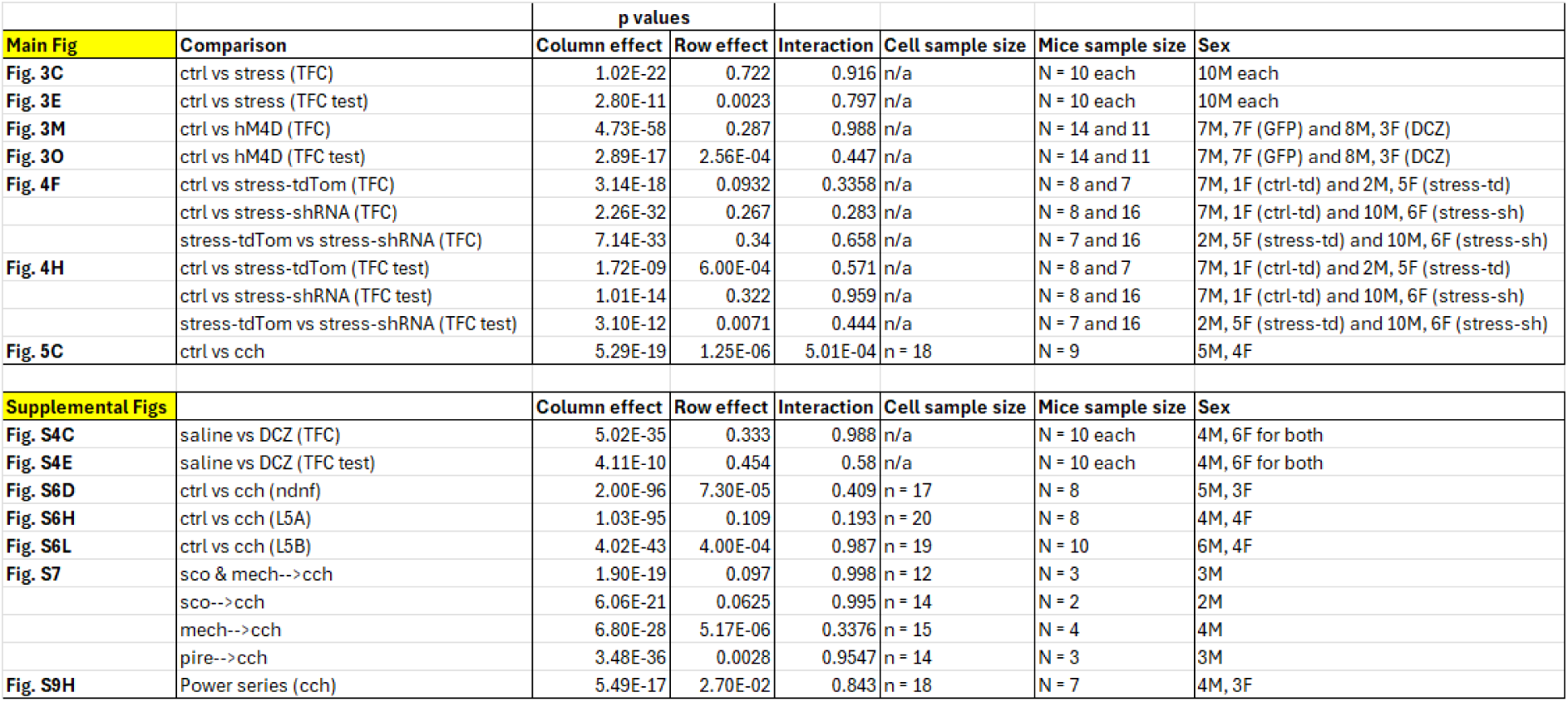
Summary of ANOVA tests and results.

**Table 3.**
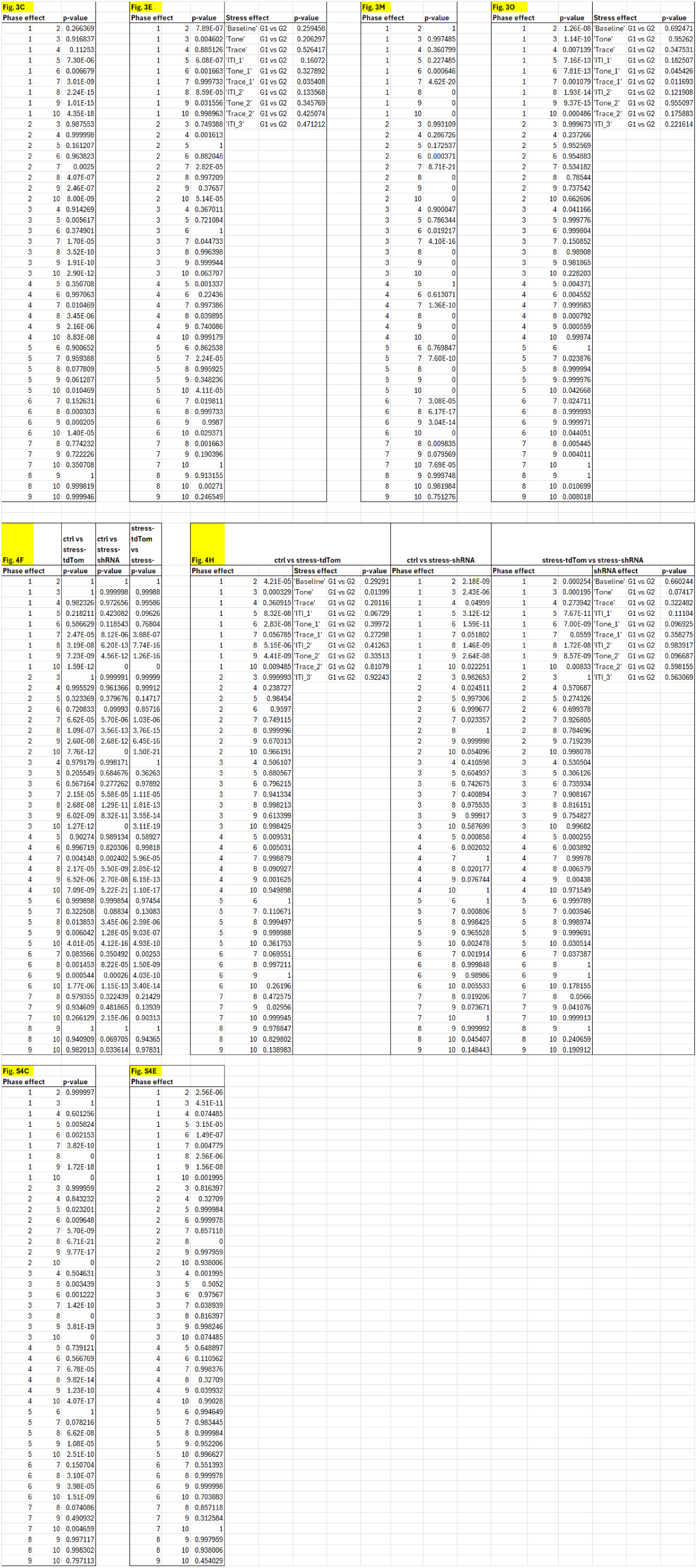
Post hoc analysis of behavioral data. Yellow highlights indicate relevant figure panels.

**Table 4.**
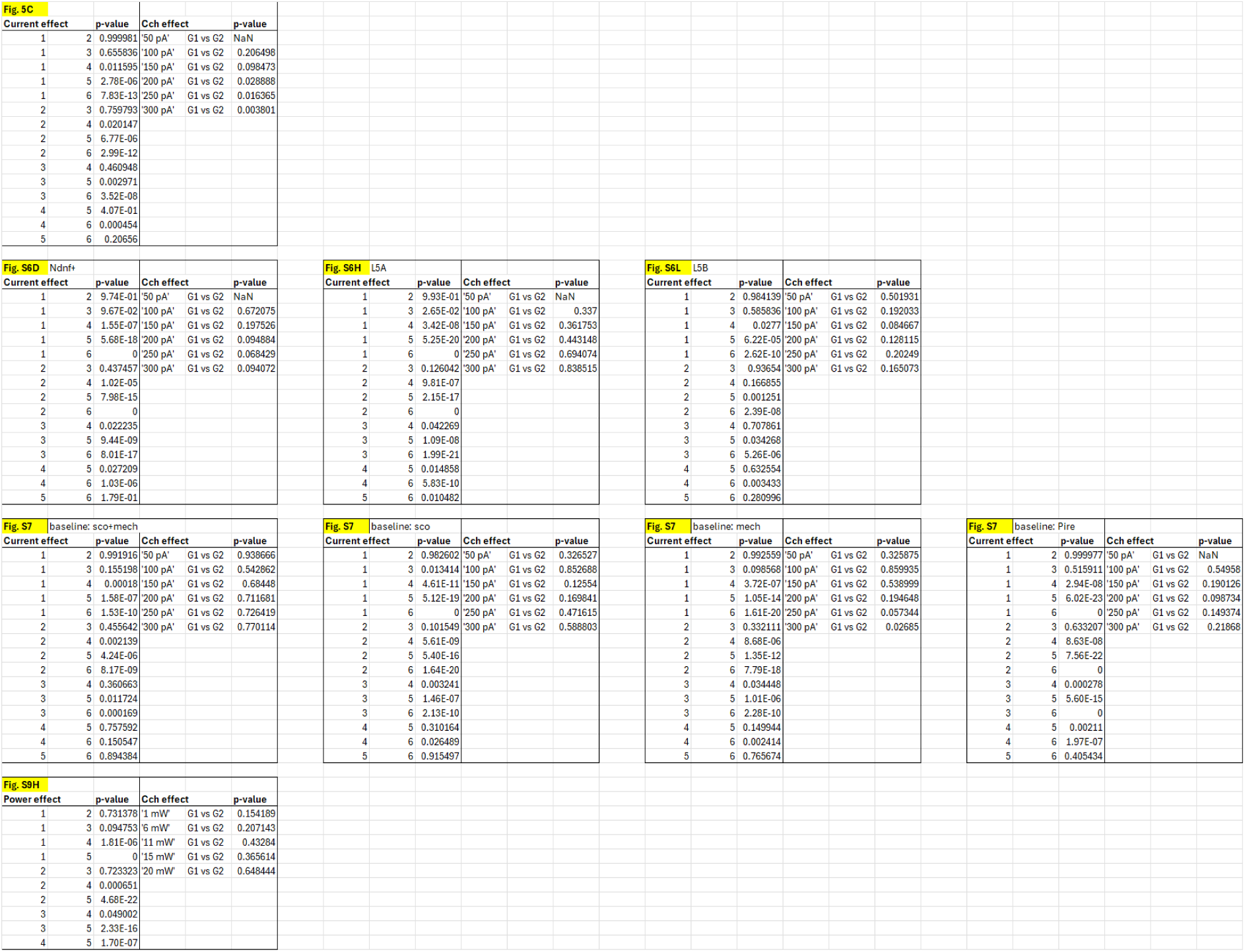
Post-hoc analysis of slice electrophysiology data. Yellow highlights indicate relevant figure panels.

## ACKNOWLEDGEMENTS

We would like to thank Kassandra Georges, Viktor Wisniewski Bendtsen, and Mie Gunni Kolmos Pedersen for supporting various aspects of the project and comments on the manuscript; Peter Bjerge, Bjarke Brix, Dennis Olesen and Matthew Peter Griffiths for technical assistance; and the Bioimaging Core Facility for the use of equipment and support. This project is supported by Lundbeck Foundation (LF) Experiment grants to N.Y. (R436-203-471) and A.T. (R436-203-471), LF Professorship to J.R. (R436-203-855), and the Danish National Research Foundation (DNRF133). H.G. is a Lundbeck Foundation Neuroscience Academy Denmark (NAD) fellow (R389-2021-1596).

## AUTHORS CONTRIBUTIONS

N.Y. conceived the study. N.Y., A.T., H.L., S.F.J., W.H.H., H.T.G., C.N.G., and T.O. conducted the experiments. N.Y., A.T., H.L., S.F.J., W.H.H., H.T.G., and C.N.G. analyzed the data. N.Y. drafted the manuscript with input from J.R., H.L., and A.T. Funding was secured by J.R., N.Y., and A.T.

---

**Figure S1.**
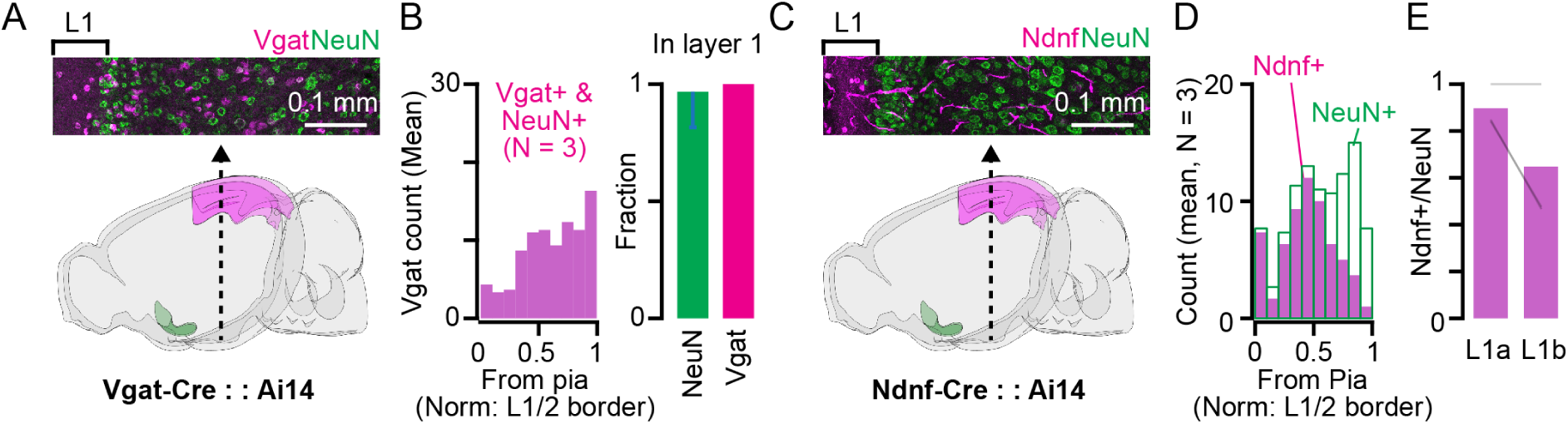
Neuronal cell types in layer 1 of the ventral retrosplenial cortex (vRSP). **(A)** A representative image of the vRSP cortical column collected from *Vgat*::Ai14 mice. The section is stained for NeuN (green), with *Vgat*^+^ neurons shown in magenta. **(B)** A histogram showing the number of *Vgat*^+^ neurons (magenta) co-labeled with NeuN (green) within vRSPL1 at different cortical depths between the pia and the border between layers 1 and 2 (N = 3). **(C)** A representative image of the vRSP cortical column collected from *Ndnf* :: Ai14 mice. The section is stained for NeuN (green), with *Ndnf*^+^ neurons shown in magenta. **(D)** A histogram showing the number of *Ndnf*^+^ neurons (magenta) co-labeled with NeuN (green) or NeuN alone at different cortical depths between the pia and the border between layers 1 and 2 (N = 3. Two lines are overlapping.). **(E)** A plot showing the number of *Ndnf^+^* neurons in the upper and lower parts of layer 1 (layer 1a and 1b). Each black line indicates data from one mouse (N = 3).

**Figure S2.**
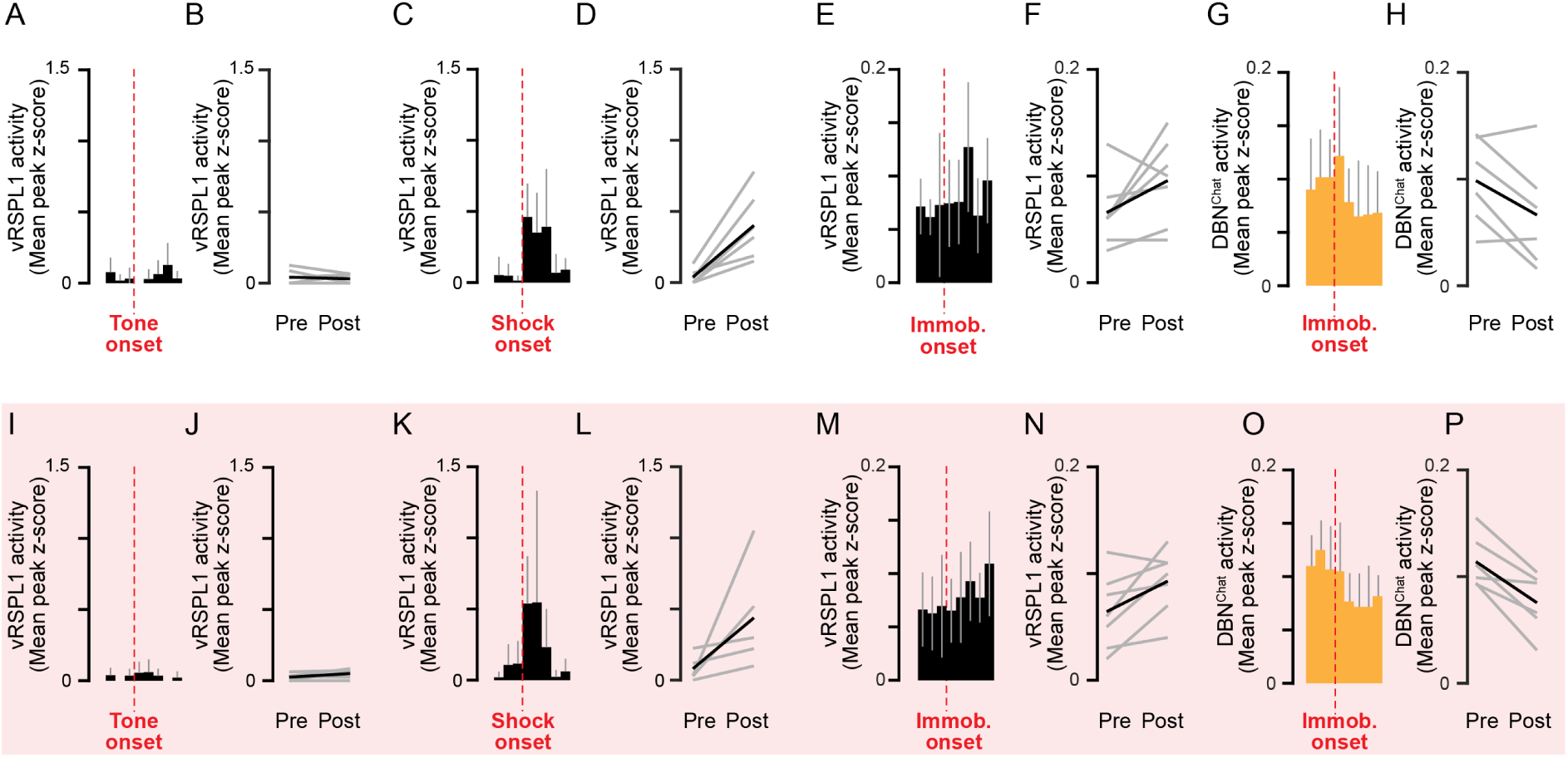
**Temporal relationship of vRSPL1 activity with respect to tone onset, footshock onset, and immobility onset.** (A) Peri-stimulus time histogram (PSTH) of vRSPL1 activity aligned to the first tone onset during TFC in control mice. Data are shown for 3 seconds before and 5 seconds after the event onset. Bin size: 1 sec. (B) Comparison of mean activity in (A) calculated over 3 second before and after the tone onset. Pre: 0.04 ± 0.05; post: 0.03 ± 0.03 (p = 0.626, paired t-test, N = 6). (C) Same as in (A), but vRSPL1 activity is aligned to the first footshock onset. (D) Same as in (B), but for data in (C). Pre: 0.04 ± 0.05; post: 0.40 ± 0.24 (p = 0.008, paired t-test, N = 6). (E) PSTH of vRSPL1 activity aligned to the immobility onset during baseline period of restraint experiment in Fig. 1M (only immobility onsets lasting more than 5 seconds and occurring at least 3 seconds after the previous immobility offset were included). (F) Comparison of mean activity calculated over 3 second before and after the immobility onset. Here, instead of using the period immediately after onset, the last 3-second bin of the 5-second window was used to calculate the mean post-immobility activity. Pre: 0.07 ± 0.03; post: 0.10 ± 0.04 (p = 0.108, paired t-test, N = 7). (G) Same as in (E), but DBN^Chat^ activity is aligned to the immobility onset. (H) Same as in (F), but for data in (G). Pre: 0.10 ± 0.04; post: 0.07 ± 0.05 (p = 0.052, paired t-test, N = 6). (I) Peri-stimulus time histogram (PSTH) of vRSPL1 activity aligned to the first tone onset during TFC in stressed mice. (J) Comparison of mean activity in (I) calculated over 3 second before and after the tone onset. Pre: 0.03 ± 0.03; post: 0.05 ± 0.04 (p = 0.155, paired t-test, N = 5). (K) Same as in (I), but vRSPL1 activity is aligned to the first footshock onset. (L) Same as in (B), but for data in (K). Pre: 0.08 ± 0.09; post: 0.44 ± 0.38 (p = 0.122, paired t-test, N = 5). (M) PSTH of vRSPL1 activity aligned to the immobility onset during post-restraint period of restraint experiment in Fig. 1M (only immobility onsets lasting more than 5 seconds and occurring at least 3 seconds after the previous immobility offset were included). (N) Same as in (B), but for data in (M). Pre: 0.06 ± 0.04; post: 0.09 ± 0.03 (p = 0.038, paired t-test, N = 7). (O) Same as in (M), but DBN^Chat^ activity is aligned to the immobility onset. (P) Same as in (N), but for data in (O). Pre: 0.11 ± 0.02; post: 0.08 ± 0.03 (p = 0.0062, paired t-test, N = 6).

**Figure S3.**
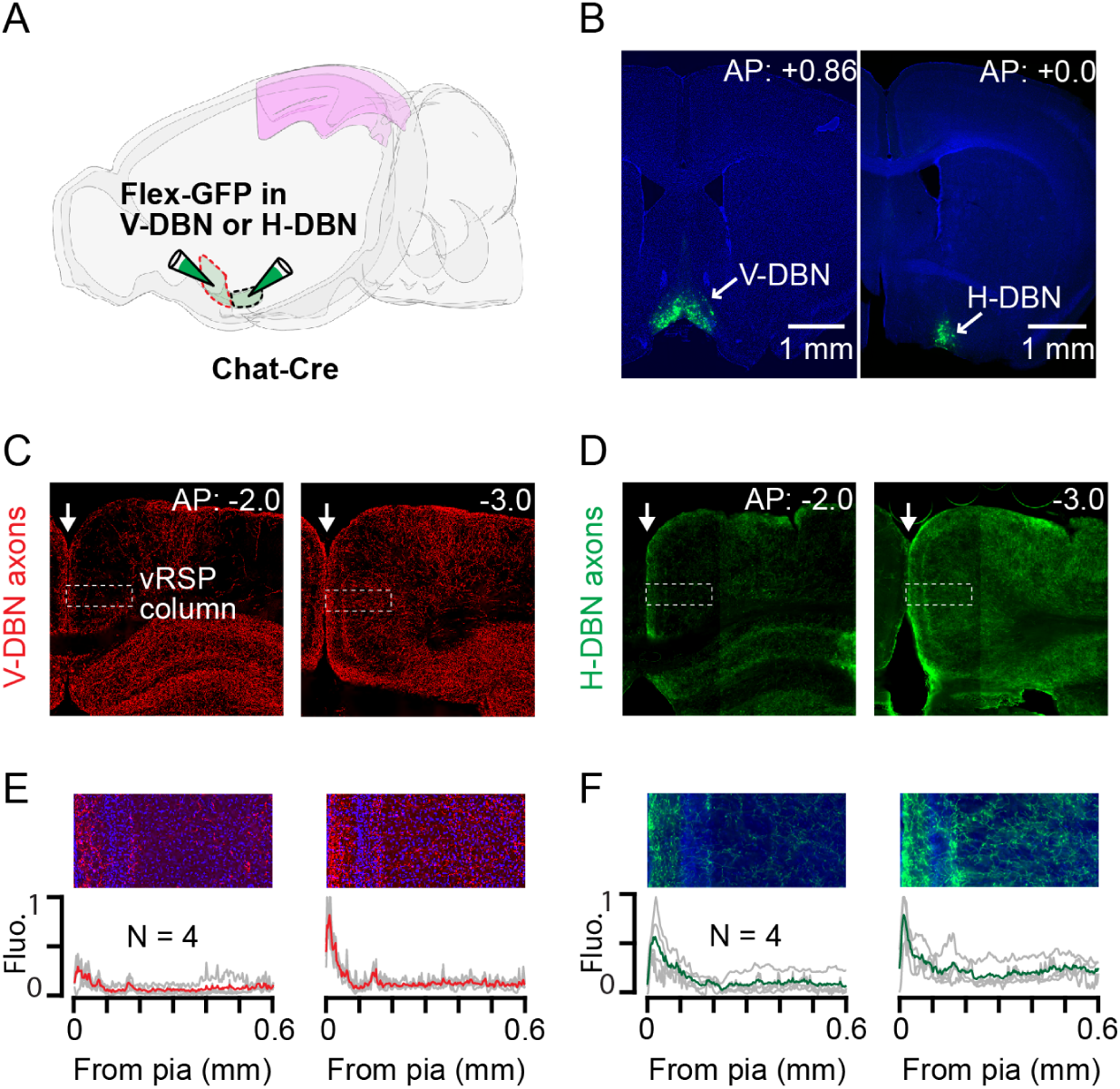
Cholinergic innervation of vRSPL1 neurons by DBN^Chat^ neurons. **(A)** Schematic of the injection performed. Each subregion of DBN (vertical DBN (V-DBN) and horizontal DBN (H-DBN)) was targeted in a separate group of mice. **(B)** Images of injection sites from two mice: one targeting the V-DBN and the other targeting the H-DBN. **(C)** Images of brain sections containing the RSP, taken at different points along the anteroposterior axis (positions indicated at the top right). Cholinergic axons from the V-DBN are shown in red. The cortical column of the vRSP is highlighted with a white rectangle. White arrow: midline. **(D)** Same as (C), but cholinergic axons from the H-DBN are shown in green. **(E)** Top: Expanded image of the vRSP cortical column from (C). Bottom: Laminar profile of the cholinergic axon fluorescent signal from the V-DBN (normalized, N = 4 mice). **(F)** Same as (E), but for the H-DBN axons.

**Figure S4.**
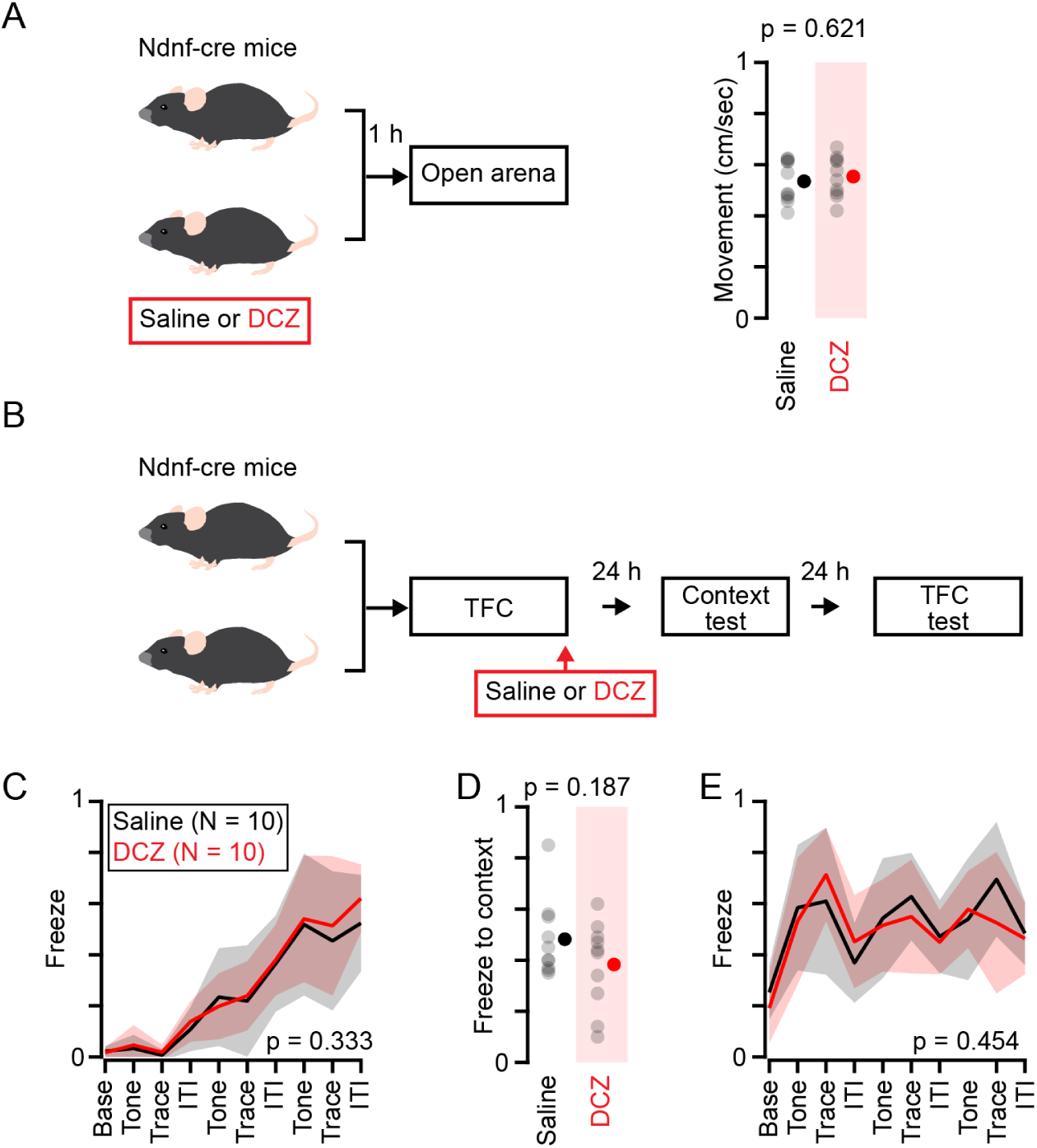
Effect of deschloroclozapine (DCZ) in control mice during the TFC protocol. **(A)** Schematic of experiment performed. One hour after i.p. injection of DCZ or saline, mice were subjected to an open area for 3 mins, where their activity was monitored (p = 0.621, unpaired t-test, N = 7 for saline and 8 for DCZ). **(B)** Schematic of experiment performed. **(C)** Freezing during TFC for control mice injected with saline (black) or DCZ (red). Two-way ANOVA: main effect of phase, F_(9,130)_ = 34.85, p = 5.02e-35; effect of group, F_(1,130)_ = 0.94, p = 0.333; interaction between phase and group, F_(9,130)_ = 0.24, p = 0.988. **(D)** Freezing during context test for saline and DCZ group (p = 0.187, unpaired t-test). **(E)** Freezing during TFC test for saline and DCZ group. Two-way ANOVA: main effect of phase, F_(9,130)_ = 8.15, p = 4.11e-10; effect of group, F_(1,130)_ = 0.56, p = 0.454; interaction between phase and group, F_(9,130)_ = 0.84, p = 0.580.

**Figure S5.**
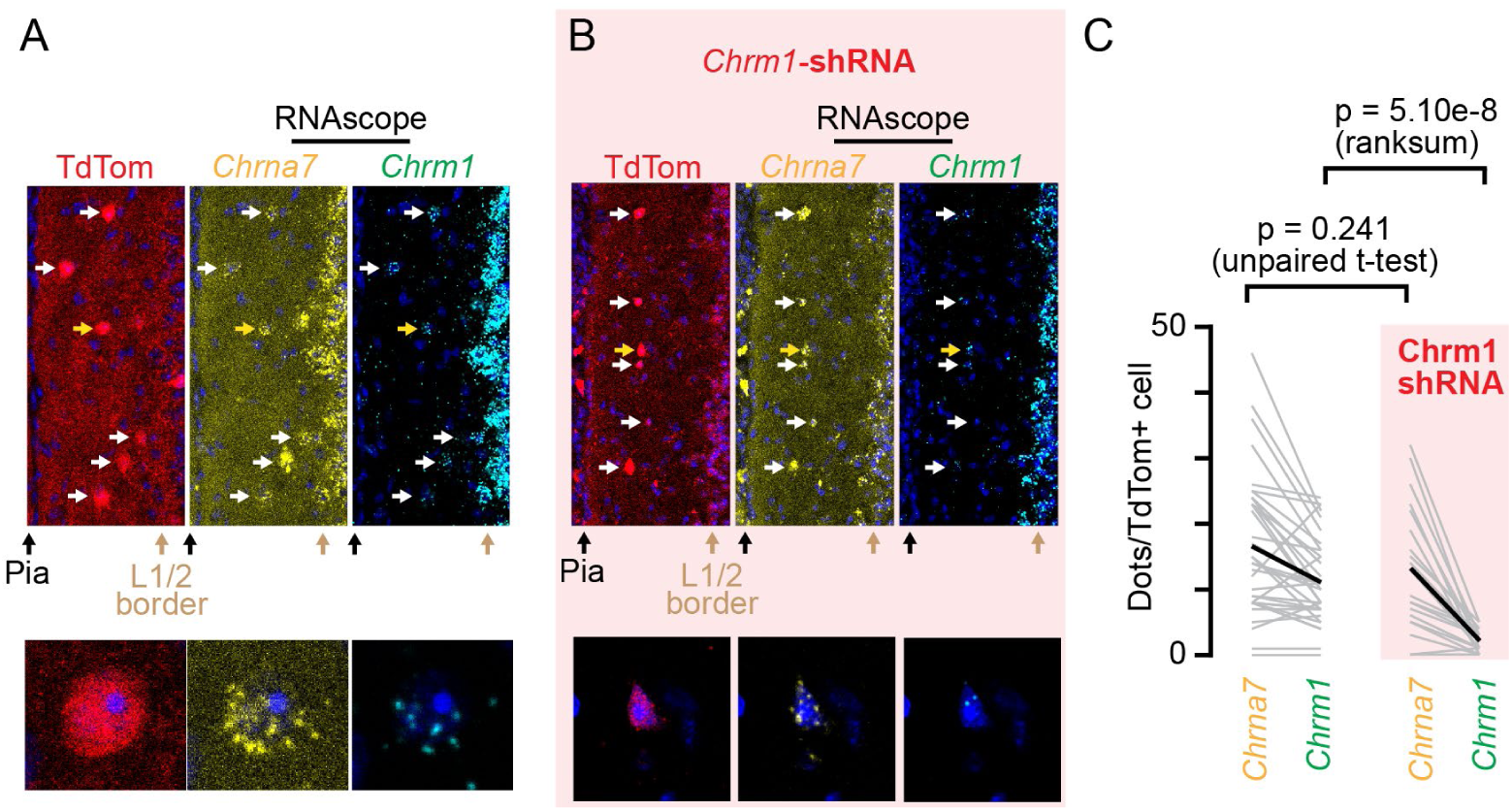
*Chrm1*-shRNA reduces *Chrm1* expression in vRSPL1 neurons. **(A)** Confocal image of a vRSPL1 section from *Vgat*::Ai14 mice, stained with RNAscope for *Chrna7* (yellow) and *Chrm1* (green). The pia and L1/2 border are indicated with black and tan arrows, respectively. White arrows mark the location of *Vgat^+^* neurons. A cell highlighted with a yellow arrow is expanded below to show individual RNA puncta. **(B)** The same as in (A) but from mice injected with Cre-dependent *Chrm1*-shRNA into vRSPL1. **(C)** Transcript abundance quantified by counting puncta per cell using s single plane (non-z-projected) image. Control mice: 11.2 ± 6.4 puncta for *Chrm1* and 16.5 ± 10.8 for *Chrna7* (p = 1.68e-4 signed-rank test, n = 35 cells, N = 3). shRNA-injected mice: 2.0 ± 1.8 for *Chrm1* and 13.0 ± 9.6 for *Chrna7* (p = 1.51e-4, signed-rank test, n = 20 cells, N = 3). Comparison between *Chrm1* expression in control and shRNA-injected mice: p = 5.10e-8 (rank-sum test). For *Chrna7*: p = 0.241 (unpaired t-test).

**Figure S6.**
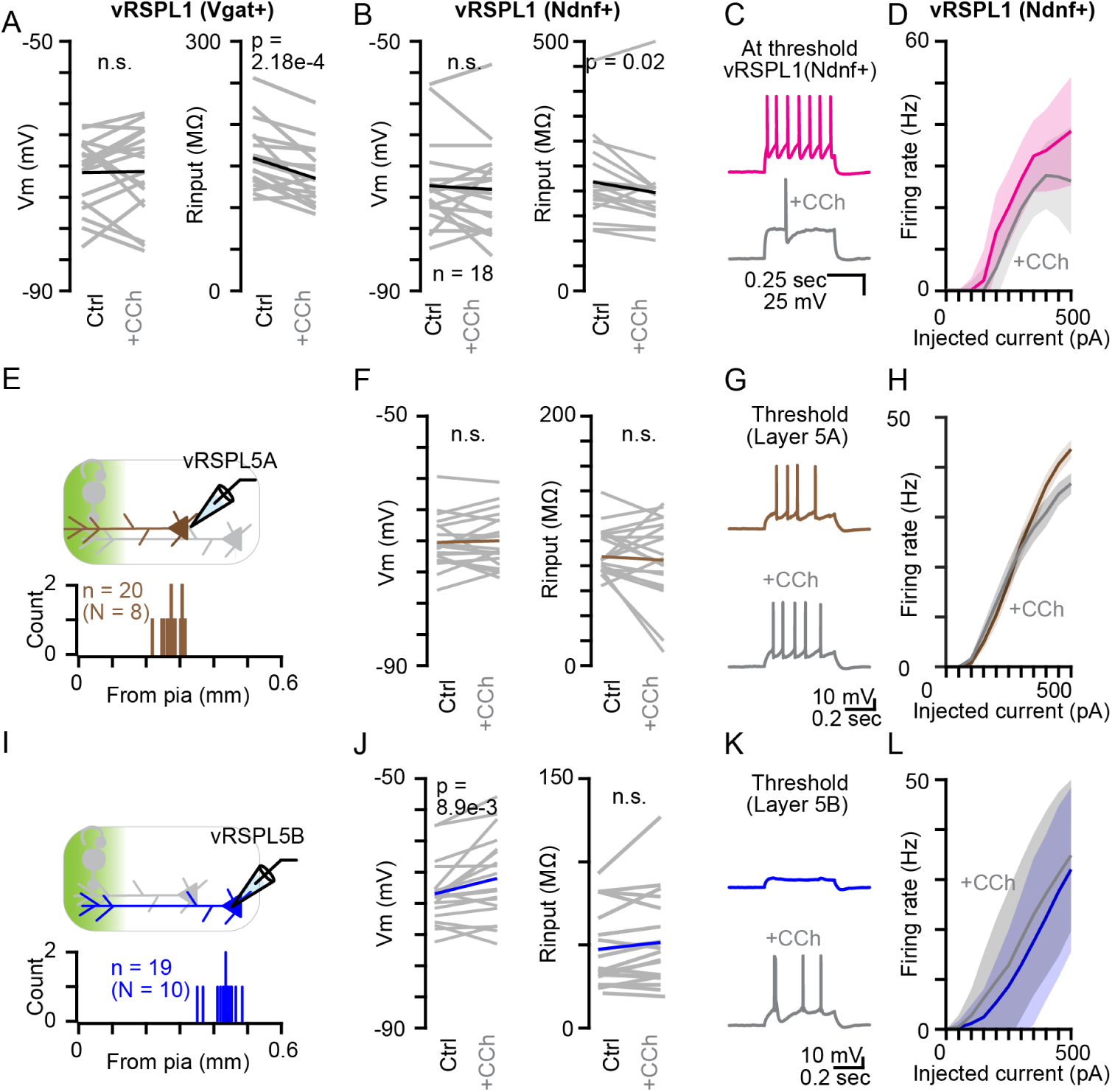
Layer-specific effects of cholinergic signaling in vRSP. **(A)** Left: Effect of CCh on the resting membrane potential (Vm) of vRSPL1 neurons identified by *Vgat^+^* labeling (*Vgat*::Ai14 mice). Vm under control and CCh conditions was −71.0 ± 5.7 mV and −70.9 ± 7.1 mV, respectively (p = 0.828, paired t-test). Right: Effect of CCh on input resistance of the same neurons, showing a reduction from 159.8 ± 39.3 MΩ to 135.2 ± 36.0 MΩ (p = 2.18e-4, paired t-test, n = 18, N = 9). **(B)** Same plot as in (A), but recording was made from vRSPL1 neurons identified by *Ndnf^+^* (using *Ndnf* :: Ai14 mice). Left: Vm under control and CCh condition was -73.2 ± 7.0 mV and -73.7 ± 7.3 mV, respectively (p = 0.569, signed-rank test). Right: Effect of CCh on input resistance of the same neurons, showing a reduction from 218.0 ± 81.8 MΩ to 196.8 ± 87.0 MΩ (p = 0.021, signed-rank test, n = 18, N = 8). **(C)** Representative spike traces from a *Ndnf^+^* vRSPL1 neuron in response to threshold current injection before (magenta) and after (gray) bath application of CCh. **(D)** Input-output (f-I) curve of *Ndnf^+^* vRSPL1 neurons before (magenta) and after (gray) CCh application. Two-way ANOVA: main effect of current, F_(14,480)_ = 60.5, p = 2.00e-96; effect of CCh, F_(1,480)_ = 16.0, p = 7.30e-5; interaction between current and CCh, F_(14,480)_ = 1.04, p = 0.409; n = 17 cells from 8 mice. **(E)** Top: Schematic of ex vivo whole-cell recordings from layer 5A pyramidal neurons in acute brain slices. Bottom: Histogram showing the distribution of cortical depths of recorded neurons. **(F)** Same as in (A), but for layer 5A neurons. Vm under control and CCh conditions was −70.2 ± 4.0 mV and −70.0 ± 4.3 mV, respectively (p = 0.682, paired t-test). Right: Effect of CCh on input resistance of the same neurons, showing a reduction from 87.4 ± 27.6 MΩ to 84.6 ± 32.5 MΩ (p = 0.546, signed-rank test, n = 20, N = 8). **(G)** Same as in (C), but from an example trace recorded from layer 5A neuron. **(H)** Same as in (D), but for data recorded from layer 5A neurons. Two-way ANOVA: main effect of current, F_(9,380)_ = 102.05, p = 1.03e-95; effect of CCh, F_(1,380)_ = 2.57, p = 0.109; interaction between current and CCh, F_(9,380)_ = 1.39, p = 0.193; n = 20 cells from 8 mice. **(I)** Top: Schematic of ex vivo whole-cell recordings from layer 5B pyramidal neurons in acute brain slices. Bottom: Histogram showing the distribution of cortical depths of recorded neurons. **(J)** Same as in (A), but for layer 5B neurons. Vm under control and CCh conditions was −68.5 ± 4.9 mV and −66.0 ± 6.8 mV, respectively (p = 0.0089, signed-rank test). Right: Effect of CCh on input resistance of the same neurons. 47.4 ± 23.0 MΩ to 51.5 ± 27.6 MΩ (p = 0.08, signed-rank test, n = 19, N = 10). **(K)** Same as in (C), but from an example trace recorded in (I). **(L)** Same as in (C), but for data recorded from layer 5B neurons. Two-way ANOVA: main effect of current, F_(9,360)_ = 33.98, p = 4.02e-43; effect of CCh, F_(1,360)_ = 12.67, p = 4.0e-4; interaction between current and CCh, F_(9,360)_ = 0.25, p = 0.987; n = 19 cells from 10 mice.

**Figure S7.**
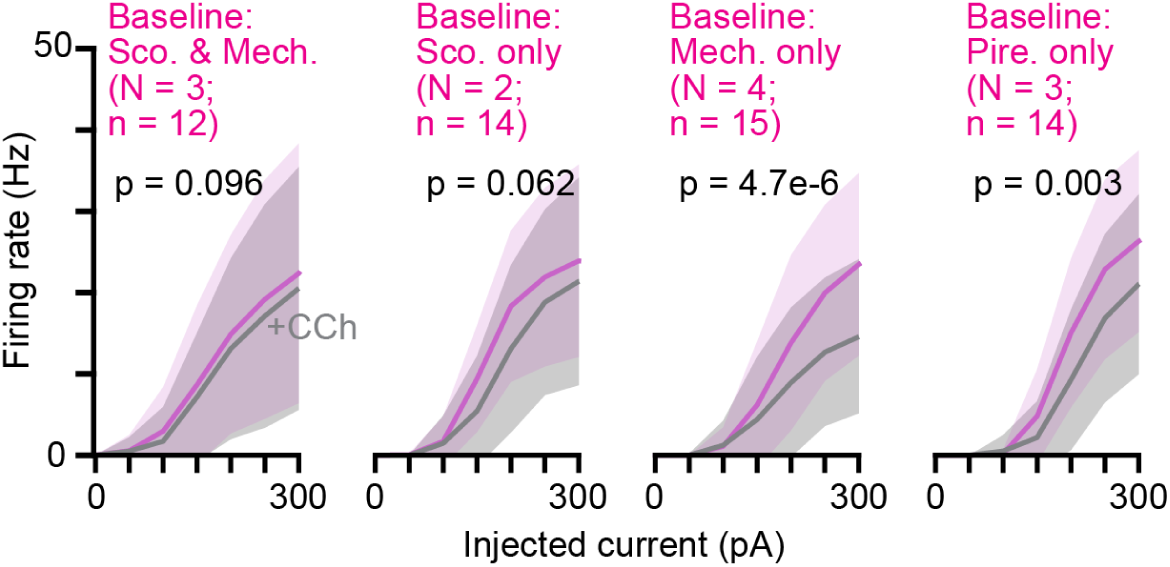
**Types of cholinergic receptors expressed in vRSPL1 neurons** Input-output (f-I) curve of vRSPL1 neurons before (magenta) and after (gray) CCh application. Different drugs were included at baseline as indicated. Two-way ANOVA: baseline with scopolamine (Sco.) and mecamylamine (Mech.): main effect of current, F_(9,220)_ = 15.46, p = 1.90e-19, effect of CCh, F_(1,220)_ = 2.78, p = 0.097; interaction between current and CCh, F_(9,220)_ = 0.14, p = 0.998; n = 12 cells from 3 mice. Baseline with Sco.: main effect of current, F_(9,260)_ = 16.12, p = 6.06e-21; effect of CCh, F_(1,260)_ = 3.5, p = 0.0625; interaction between current and CCh, F_(9,260)_ = 0.17, p = 0.995; n = 14 cells from 2 mice. Baseline with Mech.: main effect of current, F_(9,280)_ = 21.81, p = 6.80e-28; effect of CCh, F_(1,280)_ = 21.6, p = 5.17e-6; interaction between current and CCh, F_(9,280)_ = 1.14, p = 0.3376; n = 15 cells from 4 mice. Baseline with pirenzepine (Pire.): main effect of current, F_(9,260)_ = 30.64, p = 3.48e-36; effect of CCh, F_(1,260)_ = 9.14, p = 0.0028; interaction between current and CCh, F_(9,260)_ = 0.36, p = 0.9547; n = 14 cells from 3 mice.

**Figure S8.**
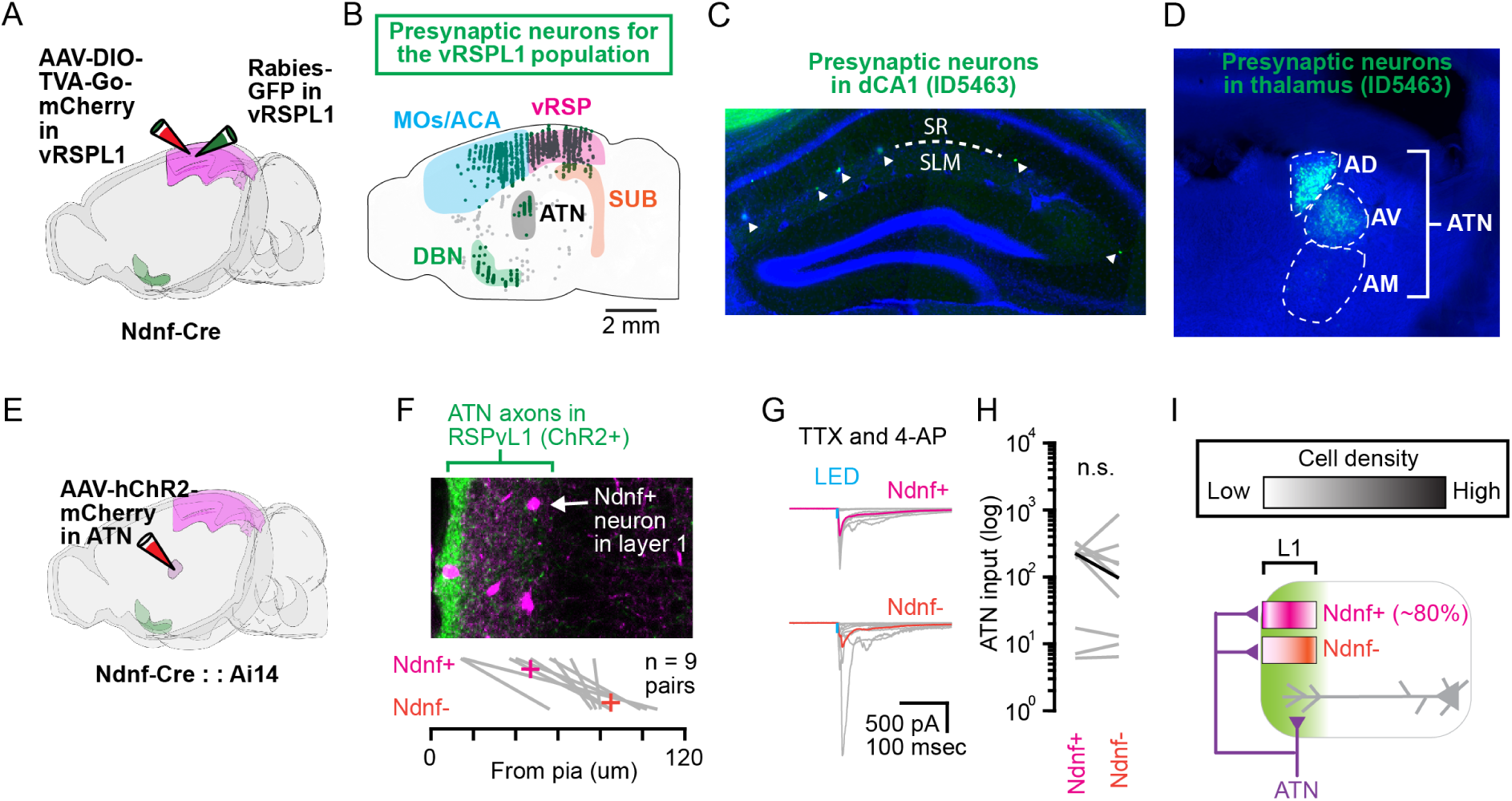
Afferent inputs to vRSPL1 neurons. **(A)** Schematic of the injection performed for monosynaptic rabies tracing. **(B)** Distribution of presynaptic neurons of vRSPL1 neurons in entire brain. A region with labeling greater than 1% of total is highlighted with the region’s name. MOs: secondary motor area. ACA: anterior cingulate area. ATN: anterior thalamic nuclei. SUB: subiculum. DBN. diagonal band nucleus. **(C)** Representative epifluorescence image of the dorsal hippocampus showing presynaptic neurons (green) of vRSPL1 neurons in this region. The border between stratum radiatum (SR) and stratum lacunosum-moleculare (SLM) is indicated with a white dashed line. Arrowheads indicate CA1–retrosplenial projection neurons. **(D)** Same as in (C), but showing an image of the thalamus containing presynaptic neurons. Thalamic subdivisions are indicated with white dashed lines. AD: anteriodorsal thalamus. AV: anteroventral thalamus. AM: anteromedial thalamus. **(E)** Schematic of the injection performed for connectivity study between ATN and vRSPL1 neurons. **(F)** Top: Confocal image of a vRSPL1 slice showing ATN axons (green) and labeled *Ndnf^+^* neurons (magenta). Bottom: Cortical depth at which the two cell types (*Ndn^f+^* (labeled) and *Ndnf^-^* (unlabeled) neurons) were sampled. Lines indicate cell pairs recorded from the same slice. **(G)** Example traces of excitatory postsynaptic currents (EPSCs) recorded from each cell pair. Each trace represents a recording from an individual cell; solid colors indicate the mean trace across all cells of each type (9 pairs from 3 mice). Recordings were made in the presence of TTX (1 µM) and 4-AP (100 µM). LED: 5 msec. **(H)** ATN input (EPSC) to each cell type. For *Ndnf^+^* neurons, −172.0 ± 129.9 pA; for *Ndnf^-^* neurons, −183.3 ± 271.5 pA (p = 0.570, signed-rank test, n = 9 pairs, N = 3). **(I)** Schematic illustrating the distribution of *Ndnf^+^* and *Ndnf^-^* neurons (based on Fig. S1) and their connectivity with ATN axons in vRSP. Green color indicates cholinergic innervation pattern found in Fig. S3.

**Figure S9.**
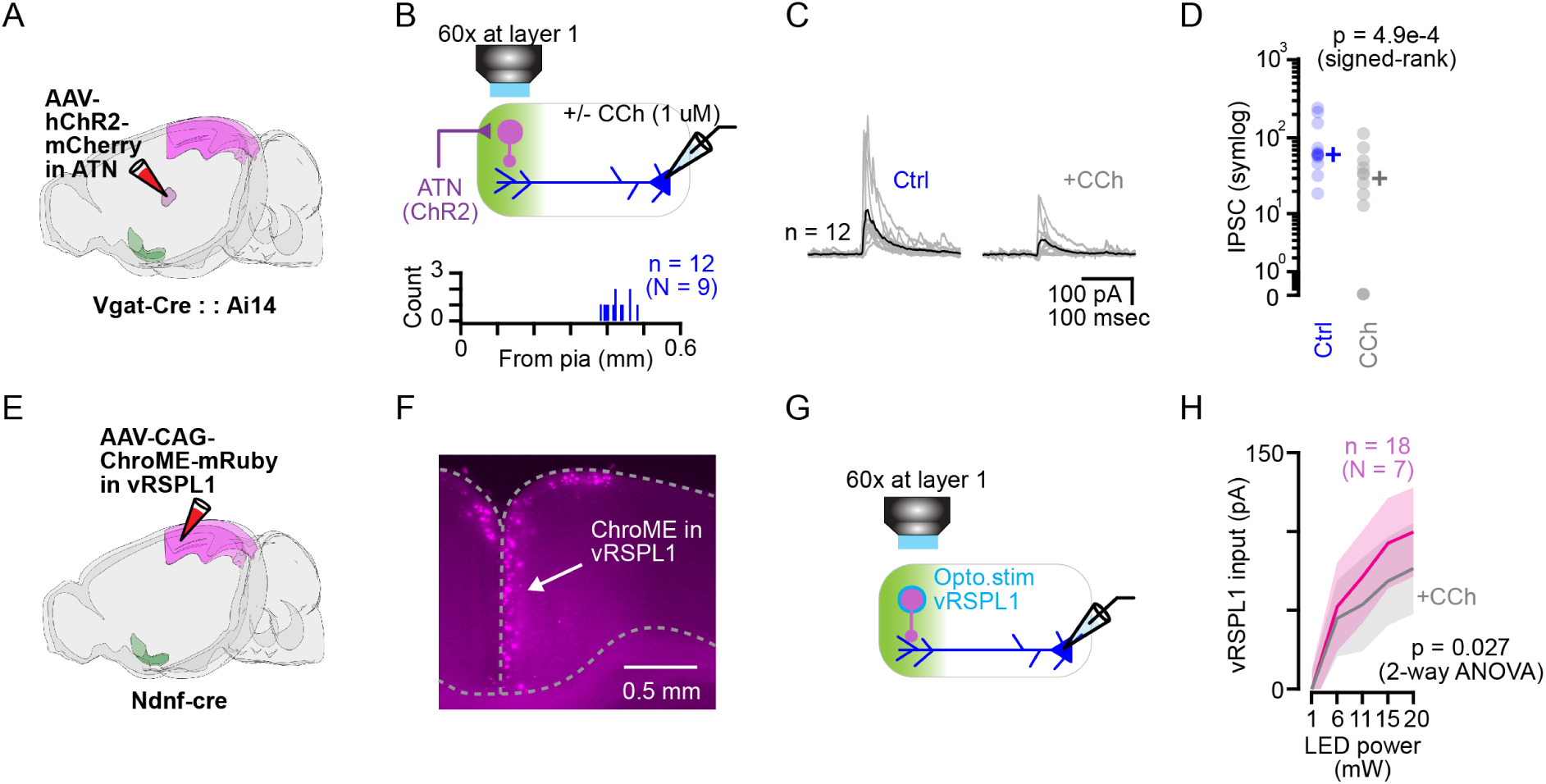
Cholinergic signaling attenuates local vRSPL1 inhibitory inputs to layer 5B pyramidal neurons. **(A)** Schematic of the injection performed. **(B)** Schematic of the experiment performed to investigate ATN-mediated feedforward inhibition onto layer 5B pyramidal neurons in vRSP. The recording site is indicated in the histogram at the bottom. **(C)** Example traces of inhibitory postsynaptic currents (IPSCs) recorded from layer 5B neurons before and after CCh application. Each trace represents a recording from an individual cell; solid colors indicate the mean trace across all cells (n = 12 cells from 9 mice). Recordings were made in drug-free condition. LED: 5 msec. **(D)** Photo-evoked IPSCs recorded before (Ctrl) and after CCh application. Ctrl: 90.4 ± 34.0 pA; CCh: 72.2 ± 33.8 pA (p = 4.88e-4, signed-rank test, n = 12 cells, N = 9 mice). **(E)** Schematic of the injection performed. **(F)** Example epifluorescence image showing expression of soma-target ChR2 (ChroME) in vRSPL1 (magenta). **(G)** Schematic of the experiment performed to investigate the effect of CCh on local inhibitory connections from vRSPL1 neurons onto layer 5B pyramidal neurons. **(H)** Inhibitory input recorded from layer 5B pyramidal neurons as a function of LED intensity before (magenta) and after (gray) CCh application. Two-way ANOVA: main effect of LED power, F_(3,172)_ = 33.21, p = 5.49e-17; effect of CCh, F_(1,172)_ = 4.98, p = 0.027; interaction between current and CCh, F_(3,172)_ = 0.28, p = 0.843; n = 18 cells from 7 mice.

